# Deep mutational scanning reveals pharmacologically relevant insights into TYK2 signaling and disease

**DOI:** 10.1101/2025.10.11.681520

**Authors:** Conor J. Howard, Nathan S. Abell, Robert R. Warneford-Thomson, Eden Mahdavi, Alan L. Su, Carmen Resnick, Nabil Mohammed, Erin M. Thompson, Emily R. Holzinger, Katrina Catalano, Abhay Hukku, Gabriel A. Mintier, Morgan MacKenzie, Bryan L. Jiang, Dora Barbosa Rabago, Angela Chan, Carolindah Ntimi, Kaitlyn N. Weiler, Stephen C. Wilson, Joseph C. Maranville, Payal R. Sheth, Robert M. Plenge, Sriram Kosuri, Diane E. Dickel

## Abstract

Tyrosine Kinase 2 (TYK2) is a genetically defined target for autoimmune disease, with first-generation inhibitors showing clinical success in some but not all associated indications. A deeper understanding of TYK2 structure-function, protein-ligand interactions, and the impact of human variants could inform next-generation therapeutics. Here, we applied Deep Mutational Scanning (DMS) to assess >23,000 amino acid substitutions across two TYK2 functions: IFN-α signaling and protein abundance. This enabled high-resolution structure-function mapping and the identification of novel allosteric sites. By coupling DMS with inhibitor treatment, we uncovered variants that modulate compound potency. We also show that human variants – both common and rare – that are protective against autoimmune phenotypes reduce TYK2 protein abundance. Together, these findings demonstrate that DMS can prospectively reveal novel druggable sites, clarify structure-activity relationships (SAR), and highlight TYK2 degradation as a potential therapeutic strategy in autoimmunity.

## Introduction

Tyrosine kinase 2 (TYK2) is a member of the Janus kinase (JAK) family of non-receptor kinases and plays an essential role in signal transduction of multiple interferon and interleukin cytokine pathways^1–3^. TYK2 binds to the intracellular domains of cytokine receptors alongside cognate JAK family members. Cytokine binding at the cell surface leads to receptor dimerization and activation of the intracellular kinases, which then phosphorylate the STAT effector transcription factors that subsequently activate downstream transcriptional programs critical for immune responses^3,4^. Human genetics studies have shown that TYK2 is a key mediator of many immune-related phenotypes. Individuals harboring rare variants that abolish TYK2 function (in homozygosity or compound heterozygosity) have immune deficiency and increased susceptibility to mycobacterial infection [OMIM: #611521]^1,2^. Additionally, genome-wide association studies (GWAS) have reproducibly identified common variants at the *TYK2* locus as associated with reduced risk to many autoimmune phenotypes, including psoriasis, rheumatoid arthritis, ulcerative colitis, multiple sclerosis, and others^5–8^. The most well studied of these protective variants (rs34536443) is a missense allele that changes the proline at residue 1104 to alanine (henceforth referred to as P1104A) that is associated with reduced activity for a subset of TYK2 signaling functions in human immune cells^8–10^. In contrast to variants that fully abolish TYK2 function, P1104A is associated with an increased risk to tuberculosis but not otherwise implicated in severe immune deficiency^8–10^. The orthologous mouse allele has been shown to protect against autoimmune encephalomyelitis in an engineered model^8^. Collectively, these findings suggested that therapies that partially, but not fully, reduce TYK2 function could be broadly effective for treating autoimmune disease with limited risk of severe infection^8^.

The selective inhibition of TYK2 is particularly appealing for the treatment of autoimmune phenotypes, as many approved therapies non-selectively inhibit multiple JAK proteins (e.g., JAK1, JAK2, and/or JAK3) and are associated with serious side effects^11^. Allosteric TYK2 inhibitors that selectively bind to the protein’s regulatory pseudokinase domain have recently emerged as promising alternatives. Deucravacitinib (Sotyktu^®^) was the first such selective TYK2 inhibitor, approved in 2022 for the treatment of moderate to severe plaque psoriasis without a black box safety warning^12–14^. Clinical trials are ongoing for this and other TYK2 inhibitors for a range of indications^15–17^. While TYK2 inhibition has shown early success in treating psoriasis, it has not demonstrated clinical efficacy in ulcerative colitis, despite the disease association with the common protective P1104A variant^18^. This underscores the need for a more comprehensive mechanistic understanding of human *TYK2* variants that protect against autoimmune disease. Such insights could inform the discovery and development of next-generation pharmaceuticals for the target.

Deep mutational scanning (DMS) is a powerful emerging method to characterize variant effects and protein structure-function relationships by using multiplexed cellular reporter assays to measure the impact of thousands of amino acid substitutions on a desired protein’s function^19–21^. In previous work, we demonstrated a DMS approach that uses cellular transcriptional assays, DNA barcoded reporters, and advanced statistical analysis methods to quantitatively and precisely measure variant effects and protein-ligand interactions for G-protein-coupled receptors (GPCRs)^22,23^. In this study, we extended these methods to a new target class (kinases) and elucidated insights into TYK2 biology that could meaningfully inform future drug discovery. In total, we assessed the effects of >99.9% of the 23,740 possible single amino acid substitutions to TYK2 on two functions: TYK2 protein abundance and immune signaling through the interferon alpha (IFN-α) pathway. By integrating abundance and signaling data, we identified protein regions with critical catalytic and allosteric functions, including novel allosteric sites with potential therapeutic value. We performed DMS assays in the presence of TYK2 inhibitors to map the function of target-drug interactions at amino acid resolution and pinpointed residues that modulate drug resistance and potency. Finally, we showed that naturally occurring TYK2 variants that reduce protein abundance are associated with protection against autoimmune disease, including rare variants with previously unknown functional or phenotypic effects. Together, these results establish a comprehensive structure–function map of TYK2, elucidate the relative importance of different drug-target interactions that could guide future compound optimization, and strongly suggest that therapies that reduce TYK2 protein abundance would be effective for the treatment of autoimmune disease.

## Results

### Effects of >23k TYK2 variants on IFN-α signaling and protein abundance

Two independent DMS assays were developed to profile the functional consequences of all 23,740 possible single missense and nonsense substitutions on two TYK2 characteristics: 1) activation of the interferon-alpha (IFN-α) immune signaling pathway, and 2) TYK2 protein abundance (**Fig. 1A**). The IFN-α DMS assay was adapted from a highly quantitative barcoded transcriptional reporter-based method we previously described^22,23^ (see **Methods** for further details). Briefly, we started with a HEK293T cell line that harbors a single-copy landing pad site for Bxb1 recombinase-mediated DNA integration of DMS reporters^23^ and knocked out all endogenous alleles of *TYK2* in this cell line using Cas9 and sequence-specific single guide RNAs (sgRNAs). We then made the resulting cell line capable of IFN-α signaling by randomly but stably integrating transgenes for two key pathway components (STAT2 and IRF9) that are not expressed in HEK293T cells. In parallel, we built a DNA reporter cassette containing *TYK2*, along with a barcoded luciferase reporter gene that is under the control of an interferon-stimulated response element (ISRE)^24^. Integrating a single copy of this reporter via Bxb1 recombination into the landing pad site of the engineered IFN-α competent HEK293T cell line resulted in reporter gene expression that was TYK2– and IFN-α dose-dependent (**Supplementary Fig. 1A**).

**Fig 1:**
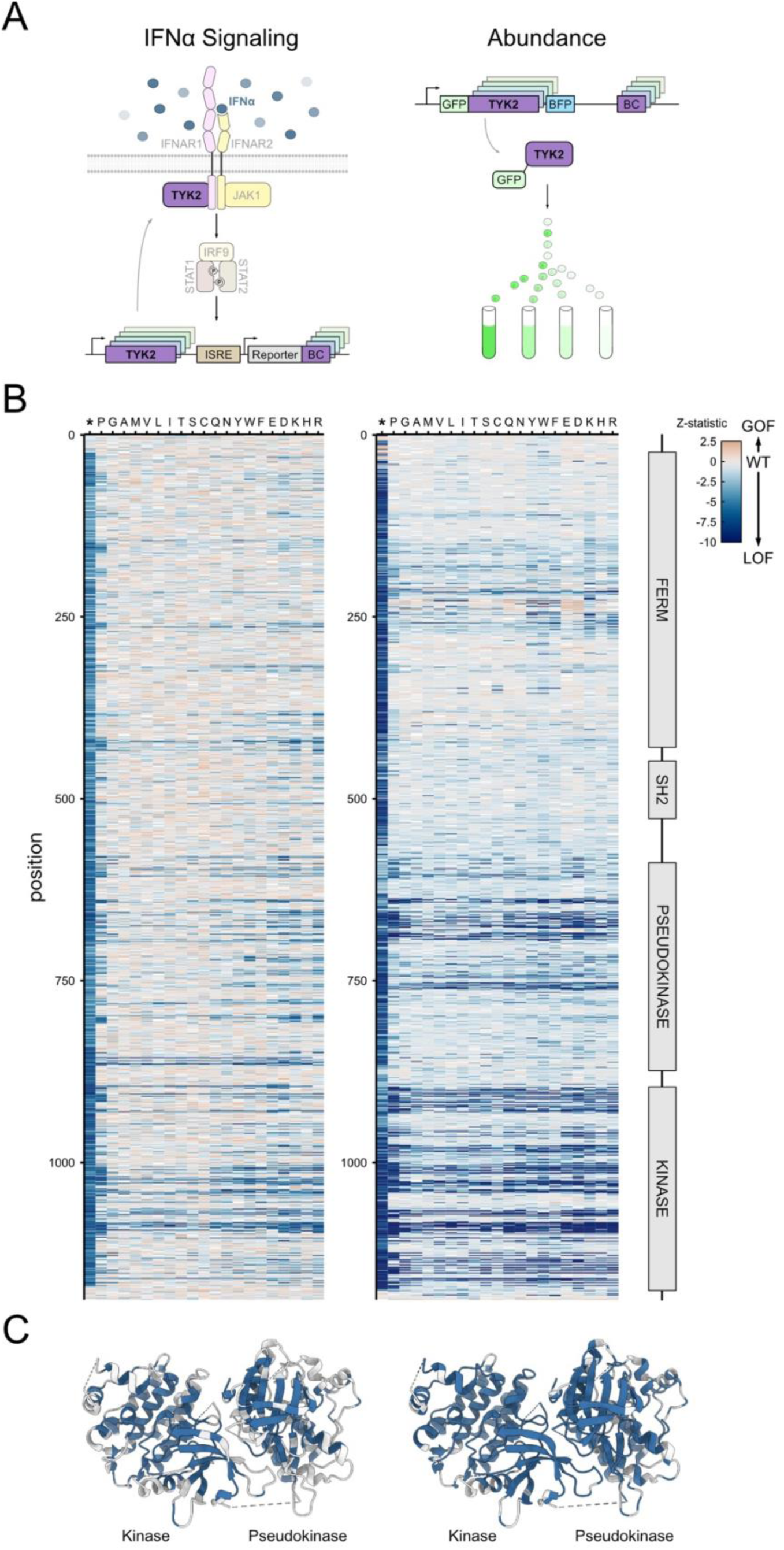
Effects of >23,000 amino acid variants on TYK2 signaling and protein abundance. **A**, Schematic showing the reporter design for TYK2 DMS assays on IFN-α signaling (left) and protein abundance (right). BC, unique DNA barcode; GFP, green fluorescent protein; ISRE, interferon-stimulated response element. **B**, Heatmaps showing the functional effects of each TYK2 variant, represented as Z-scores, for IFN-α signaling (left, 100 U/mL IFN-α condition) and protein abundance (right). Rows and columns of the heatmap correspond to amino acid position and variant identity, respectively. TYK2 protein domains are shown on the far right. GOF, gain-of-function; LOF, loss-of-function; WT, wild-type. **C**, Structures of the TYK2 kinase and pseudokinase domains (PDB: 4OLI) with residues colored blue to indicate positions that have at least two significant LOF variants at that position in the IFN-α signaling (left) or protein abundance (right) assay (FDR < 0.01).

For the TYK2 protein abundance assay, we used the same engineered cell line described above and adapted the Variant Abundance by Massively Parallel sequencing (VAMP-seq) reporter method^25^ (see **Methods** for further details). Briefly, green fluorescent protein (GFP) was appended to the amino-terminus of TYK2 (**Fig. 1A**), and this was linked to a second reporter gene (blue fluorescent protein, BFP) that served as a transcriptional control. Fluorescence-activated cell sorting (FACS) of GFP normalized to BFP clearly distinguished wild-type TYK2 from variants (e.g., E154X, K930E) that reduce TYK2 stability (**Supplementary Fig. 1B**).

Finally, we previously described the development of a negative binomial mixed effect model-based analysis method for DMS data^23^, which we used for the IFN-α assay. Existing VAMP-seq analysis methods do not harness the full statistical power of the DNA barcodes present in the reporter design. Therefore, we developed a new analysis framework for this assay (described in detail in **Methods**). Briefly, we used a negative binomial generalized linear mixed model to explicitly account for both covariates and the high degree of barcode multiplexing, while allowing inference of mean shifts between conditions. We applied the results of this model to define and compute secondary quantities of interest, like contrasts across conditions and bin midpoints for VAMP-seq. This prevents loss of information induced by aggregating barcodes linked to the same variant, by treating them as independent experiments in distinct cells. With these tools in hand, we then built comprehensive DMS libraries and assessed the effects of all possible single amino acid substitutions on TYK2 functions.

For both assays, we generated high-quality DMS libraries with near complete (>99.9%) variant coverage. Of 23,740 possible single missense or nonsense variants, the IFN-α and abundance libraries included 23,737 and 23,716 variants, respectively (**Fig. 1B**). Each variant was uniquely represented many times in both libraries, with a median of 25 and 15 unique barcodes per variant for the IFN-α and abundance DMS libraries, respectively. These factors, combined with running the IFN-α DMS assay under a range of cytokine stimulation conditions (**Table 1**), allowed us to accurately and comprehensively assess quantitative variant effects for nearly all possible single amino acid substitutions and nonsense variants of TYK2 (**Fig. 1B, Supplementary Figs. 2, 3A,B**). For example, running the IFN-α DMS assay under high stimulation (100 U/mL IFN-α) identified 2,578 loss-of-function (LOF) variants (∼11% of tested variants, FDR < 0.05, **Supplementary Fig. 3A**). For the protein abundance assay, 4,718 variants (∼20%) significantly reduced protein abundance (FDR < 0.05, **Supplementary Fig. 3A**).

**Table 1:** Experimental conditions assayed by DMS.

| Reporter | Cytokine Stimulation | Inhibitor |
| --- | --- | --- |
| IFN-α | -- | -- |
| IFN-α | 1 U/mL IFN-α | -- |
| IFN-α | 10 U/mL IFN-α | -- |
| IFN-α | 100 U/mL IFN-α | -- |
| IFN-α | 100 U/mL IFN-α | 20 nM BMS-986202 (IC<sub>75</sub>) |
| IFN-α | 100 U/mL IFN-α | 1 μM BMS-986202 (IC<sub>99</sub>) |
| IFN-α | 100 U/mL IFN-α | 10 μM BMS-986202 (>IC<sub>99</sub>) |
| IFN-α | 100 U/mL IFN-α | 7 nM NDI-034858 (IC<sub>75</sub>) |
| IFN-α | 100 U/mL IFN-α | 1 μM NDI-034858 (IC<sub>99</sub>) |
| Abundance | -- | -- |

IC: inhibitory concentration

Benchmarking these results against orthogonal structural data and variant classifications supports the high quality of these DMS assays. As expected, variants that are LOF in either assay cluster in the cores of the major domains of the protein (**Fig. 1C**). Furthermore, variants predicted to be pathogenic by AlphaMissense^26^ have significantly lower IFN-α activity and TYK2 protein abundance scores than those predicted to be benign (**Supplementary Fig. 3C**; e.g., IFN-α assay median z-scores were –1.1 versus –0.1 for variants predicted by AlphaMissense to be pathogenic versus benign, respectively; p < 2.2e-16, Kolmogorov-Smirnov test). From literature searches, we identified a panel of 32 TYK2 variants that were previously characterized, including 15 reported to be LOF^1,2,9,27–37^. Of these, 11/15 (73.4%) were identified as significantly loss-of-function in the IFN-α assay (100 U/mL stimulation), with the same number LOF in the protein abundance assay (**Supplementary Fig. 3D**). Finally, although the majority of *TYK2* variants in ClinVar have unknown functional or clinical significance, our DMS results are consistent with those that do have reported classifications. Specifically, only three variants are reported as pathogenic in ClinVar, 19 are classified as benign or likely benign, and 391 are of uncertain significance^38^. For the IFN-α signaling assay, 2/3 (66.7%) ClinVar pathogenic variants are significantly LOF and 18/18 (100%) ClinVar benign/likely benign variants show no significant effect on function (**Supplementary Fig. 3E**). Results were similar for the protein abundance assay (**Supplementary Fig. 3E**, 66.7% of ClinVar pathogenic variants show significantly reduced abundance, 77.8% of ClinVar benign variants show no significant effect on abundance). Collectively, these results demonstrate the high quality and broad coverage of these deep scans of TYK2 function.

Such highly quantitative DMS results can, in principle, be used to glean mechanistic and pharmacological insights to inform decision-making during drug discovery and development^23,39^. For example, these methods, when performed on multiple protein functions and combined with varying assay conditions, should enable prospective identification of distinct allosteric sites on the target, predict variant-specific drug responses or potential resistance mechanisms, assess the relative functions of target-ligand interactions, and identify the mechanistic effects of human variants to guide modality selection for a therapeutic program. To this end, we performed the IFN-α signaling assay with a variety of TYK2 inhibitor conditions (**Table 1**), generating a total of 356,089 estimates of TYK2 variant-condition effects, and integrated together results from both functional assays (cytokine signaling and protein abundance) to build an in-depth understanding of TYK2 to enable future drug discovery efforts.

### Identifying key functional sites of TYK2

JAK kinases bind the intracellular tail of cytokine receptors, and, in their inactive state, the kinase and pseudokinase domains are tightly bound to one another. Upon cytokine binding, the receptors dimerize and bring together their cognate kinases, leading to activation of the kinase domains^40^, likely through an unfurling of the kinase domain from the pseudokinase domain^41^. A growing number of TYK2 inhibitors, including the recently approved therapy deucravacitinib (Sotyktu^®^), target a known allosteric region within the pseudokinase domain of TYK2 and harnesses the regulatory function of this domain to inhibit TYK2’s kinase activity^42,43^. Targeting this allosteric site affords better target specificity, limiting off-target interactions of the drug with other members of the JAK family and reducing the risk of severe side-effects^15,44^. The identification of other key allosteric sites would provide additional desirable nodes for therapeutic modulation of TYK2. By systematically mapping protein function to structure, DMS is a potentially powerful tool for this purpose^39^. Specific to the identification of allosteric sites, the IFN-α signaling assay identifies variants that impact TYK2 activity by <u>any</u> mechanism (e.g., inactivating catalytic activity, reducing protein expression, etc.), while the protein abundance assay specifically identifies variants that impact protein abundance (e.g., via folding, stability, trafficking, and/or turnover). Therefore, we hypothesized that intersecting the cytokine signaling and abundance DMS results to define variants that impact the cytokine signaling function without also affecting protein abundance could identify key allosteric and catalytic domains within TYK2 (**Fig. 2A**).

**Fig. 2:**
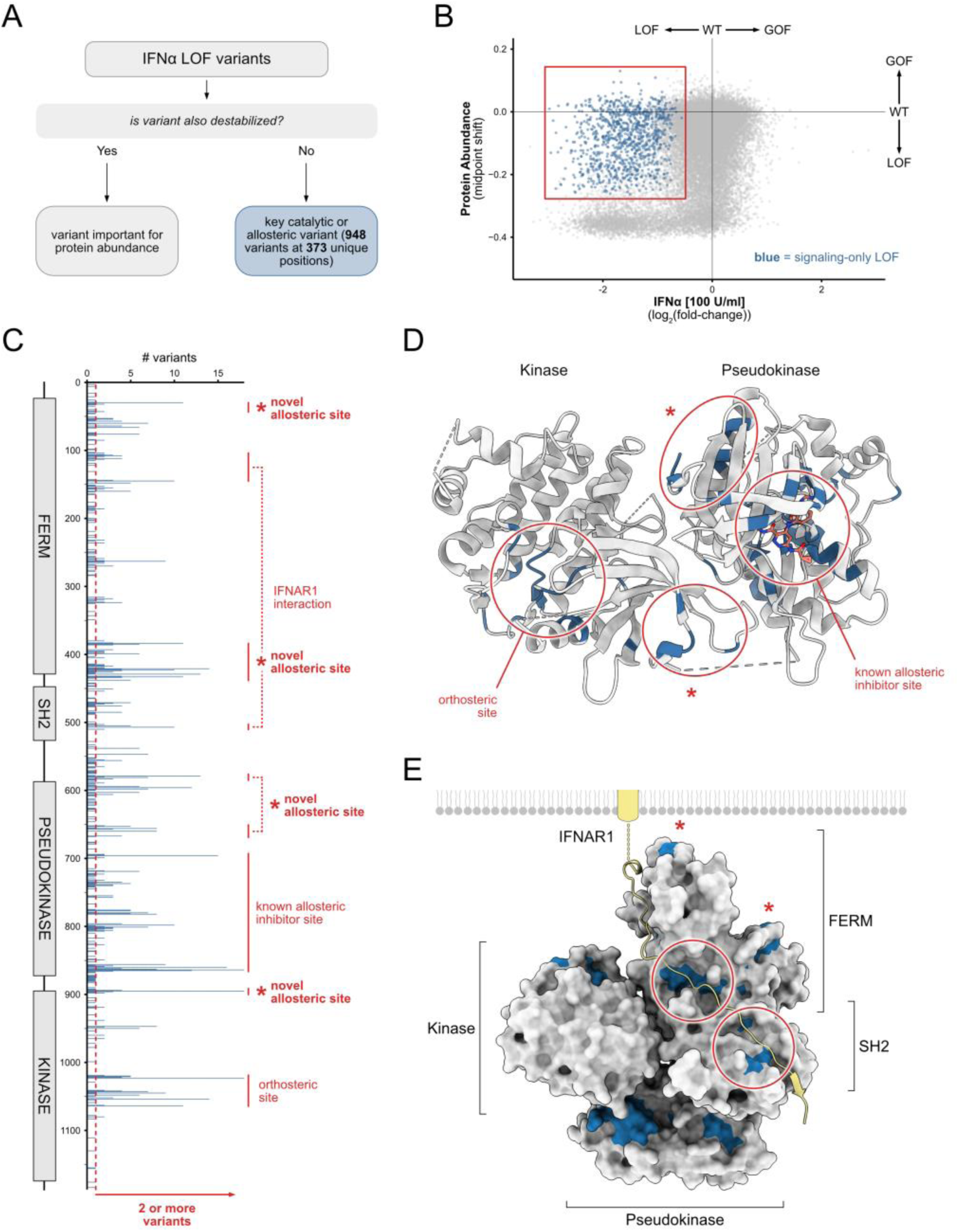
Identification of allosteric and other functionally important sites of TYK2. **A**, Schematic showing approach to identify key functional sites by pinpointing TYK2 variants that impact IFN-α signaling but not protein abundance. LOF, loss-of-function. **B**, Scatterplot of individual TYK2 variant effects on IFN-α signaling activity (x-axis) and TYK2 protein abundance (y-axis). Variants highlighted in blue were classified as signaling-only LOF (see **Methods** for details). GOF, gain-of-function; WT, wild-type. **C**, Count of signaling-only LOF variants (horizontal axis) at each residue of TYK2 (vertical axis), shown alongside the functional domains of the protein (left). Regions important for TYK2 function are highlighted in red, including known catalytic, known allosteric, and novel allosteric sites predicted by this analysis. **D**, Structure of the TYK2 kinase and pseudokinase domains (PDB: 4OLI) with amino acid positions that have at least two significant signaling-only LOF variants colored blue. Novel allosteric sites predicted by this analysis are labeled with an asterisk (*). **E**, Structural model of inactive full-length TYK2 (colored as in **D**) bound to the intracellular tail of the IFNAR1 receptor (yellow). See **Methods** for details on how this composite model was constructed.

**Fig. 3:**
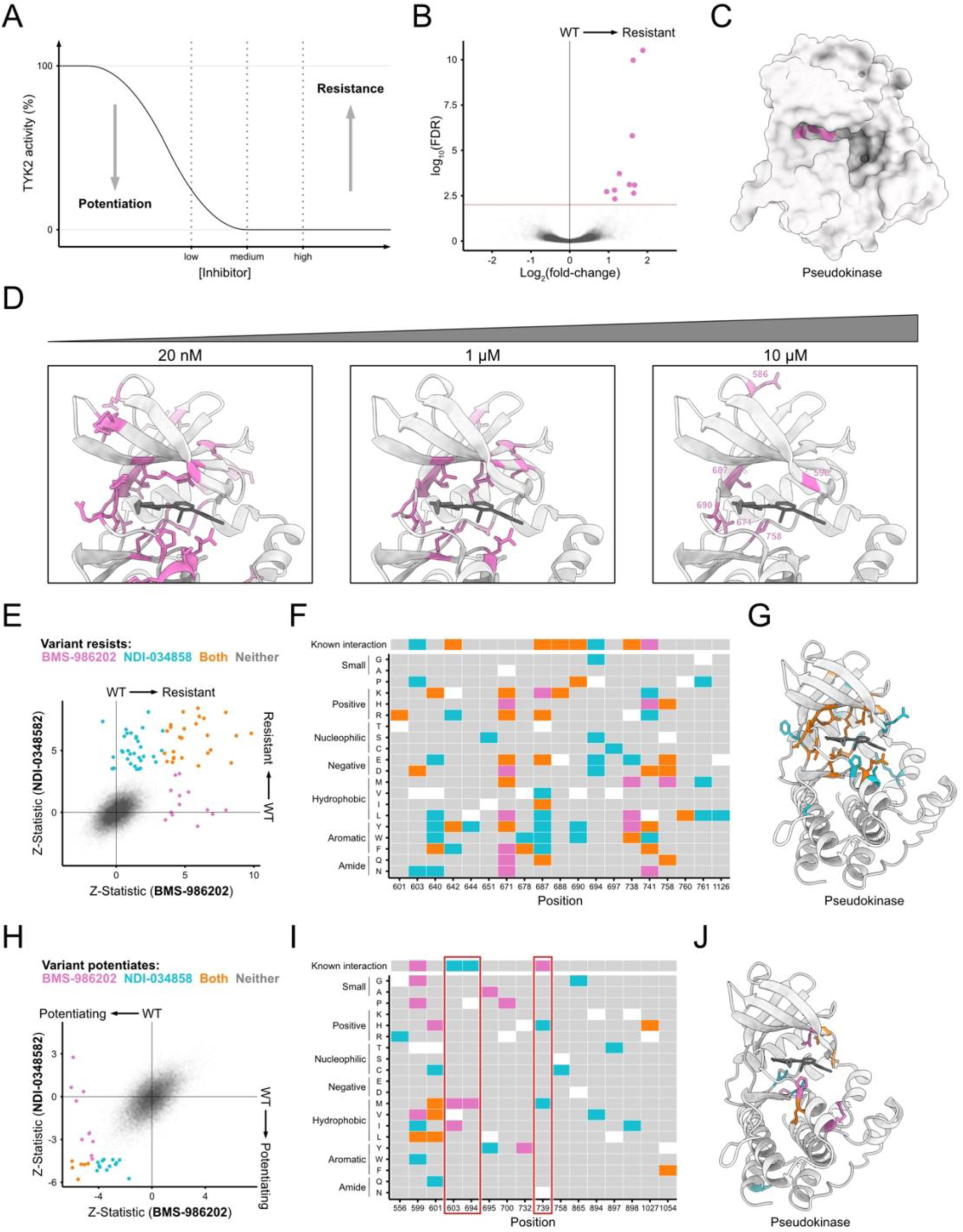
Functional characterization of protein-drug interactions. **A**, Conceptual schematic of how inhibitor dose influences variant effect interpretation. **B**, Volcano plot of variant effects under very high dose (>IC_99_) of BMS-986202, with significant drug-resistant variants colored pink (FDR < 0.01 used as significance threshold throughout figure). **C**, Surface representation of the TYK2 pseudokinase domain (PDB: 6NZP) colored as in **B**. **D**, Allosteric binding site of the TYK2 pseudokinase domain with inhibitor BMS-986202 bound (dark gray) and positions with significant drug-resistance colored pink (as in C) for increasing concentrations of BMS-986202. **E**, Scatter plot of variant effects (Z-statistic) in the presence of inhibitor (IC_99_ concentration), with variants colored by whether they exhibit resistance to BMS-986202 (pink), NDI-034858 (cyan), or both (orange). **F**, Heatmap of drug-resistant positions (x-axis) and variants (y-axis) colored as in **E**. **G**, Structure (PDB: 6NZP) of TYK2 pseudokinase domain (colored as in **E**). **H, I, J**, Variants that exhibit drug-potentiation to each inhibitor (at IC_75_ concentrations), displayed as in **E**, **F**, and **G**, respectively.

To this end, we partitioned variants that were LOF in the IFN-α assay (100 U/mL stimulation condition) by whether they also impacted TYK2 abundance (see **Methods** for details). This identified a subset of 948 variants that lead to defects in signaling but not abundance, spread across 373 positions (**Fig. 2A-C, Supplementary Fig. 4, Table S1**). Many positions harbor just one such variant, and these are broadly dispersed throughout the protein. To identify key functional sites, we focused on those positions where there are two or more variants that impact signaling but not abundance (**Fig. 2C, Supplementary Fig. 4**). This resulted in 162 positions, including several where most amino acid substitutions at the given position impacted cytokine signaling but not protein abundance (**Fig. 2C**). Importantly, these positions include three key catalytic residues in the orthosteric site of the protein (E947, D1023, D1041), in addition to non-catalytic residues in the orthosteric pocket (Positions 1021, 1022, 1063, 1064) and the activation segment (Positions 1042, 1043, 1045, 1048, 1052, 1054) that are known to regulate kinase activity^45^ (**Fig. 2C,D**). This also highlighted multiple residues in the known allosteric binding pocket of TYK2 (Positions 594, 596, 598, 603, 606, 694, 695, 696, 737, 739, 740, 757)^15,46^ (**Fig. 2C,D**). Collectively, this confirms that the approach can identify key sites of functional regulation of TYK2.

In addition to key sites of known function, we capture intriguing, less well-characterized functional positions. Two distinct regions at the kinase-pseudokinase domain interface show clusters of positions where mutation impacts cytokine signaling but not abundance (**Fig. 2C,D**, **Supplementary Fig. 4**), suggesting that these residues are important for the regulation of the kinase domain by the pseudokinase domain^33^. We also identified intriguing potential allosteric regions within the FERM and SH2 domains of the protein, which are less well-characterized than the kinase and pseudokinase domains (**Fig. 2C**). Because a full-length structure of TYK2 does not currently exist, we used available partial structures and a full-length structure of a related protein (JAK1) to construct a full-length model of TYK2 bound to a cytokine receptor and mapped these sites onto it (**Fig. 2E**, see **Methods** for details). This revealed two surfaces that interface with the tail of the interferon alpha and beta receptor subunit 1 (IFNAR1) (**Fig. 2E**, circled), including several residues known to interact directly with IFNAR1 (L144, S476, T477 and R503)^47^. Our data also captured two clusters of residues along the FERM domain surface that faces the plasma membrane (**Fig. 2E**, asterisks), suggesting novel target sites that regulate TYK2 activity. In summary, our complementary signaling and abundance DMS datasets comprehensively map key functional sites of TYK2. In addition to known catalytic and allosteric positions, this includes novel allosteric sites that may modulate intra– and inter-protein interactions that could be explored for future therapeutic targeting of TYK2.

### Functional insights into protein-drug interactions

In addition to providing key insights into protein function, DMS can be used to elucidate functional protein-drug interactions. This includes identifying variants that are likely to lead to drug resistance and, with sufficiently sensitive methods, those that potentiate (i.e., improve) the effects of therapies^23,48–52^. In principle, variants that improve the effects of drug candidates could be harnessed in structure activity relationship (SAR)-based methods to guide the design of more potent compounds^23^. To gain additional understanding of how selective allosteric inhibitors of TYK2 interact with its pseudokinase domain – and how these interactions may be improved – we performed IFN-α signaling DMS experiments in the presence of two inhibitor compounds: BMS-986202^53^ and NDI-034858 (alias: TAK-279)^46^ (**Supplementary Fig. 5A**). We confirmed that these two compounds result in dose-dependent inhibition of wild-type TYK2’s IFN-α signaling activity in our assay (**Supplementary Fig. 5B**). We then ran the IFN-α DMS assay under a range of inhibitor concentrations to probe two distinct questions: 1) to identify positions that are critical for inhibitor activity, we looked for variants that retain TYK2 activity under *high* drug concentrations (i.e., drug-resistant variants); and 2) to identify variants that improve inhibitor activity (i.e., drug-potentiating variants) we identified variants that led to further inhibition of TYK2 activity under *sub-maximal* drug concentrations (**Table 1**, **Fig. 3A**).

First, we assessed variant resistance to a range of high doses of the first inhibitor compound (BMS-986202). At a very high dose of BMS-986202 (10 uM, well above the 99% inhibitory concentration [>IC_99_]) we identified ten drug-resistant variants located at five unique positions (**Fig. 3B,C**, right panel of **D**; significance of FDR < 0.01). Many of these variants occur at residues that have been shown through crystallography to be important for protein-drug interactions, validating the DMS prediction that these positions are critical for direct binding and/or steric accommodation of the inhibitor. For example, position 690 forms a hydrogen bond interaction between TYK2 and compounds structurally similar to BMS-986202^54^. Changing this position from valine to proline results in resistance to BMS-986202. Additionally, residue A671 creates an open “alanine pocket” that was exploited in the design of BMS-986202 to confer target specificity^54^, and changing it to E, K, L, Q, or Y leads to drug resistance. Three of the remaining four resistance variants (T687R, S758D, and S758H) occur at other proximal positions within the binding pocket.

Using a high dose of inhibitor exposes residues that are critical for compound binding, and we reasoned that exploring lower doses would reveal residues that make more subtle, quantitative contributions to protein-drug interactions. Indeed, as inhibitor concentrations were lowered (to IC_99_ and IC_75_), we observed an expansion of the spatial distribution of drug-resistant variants (**Fig. 3D**). Lower doses revealed drug-resistant variants at additional positions within the binding pocket, suggesting that these positions more weakly contribute to the binding and activity of the drug. We also observed drug-resistant variants at lower doses that are within the pseudokinase domain but are well outside of the binding pocket (**Fig. 3D**). These more distal positions likely represent residues important for the allosteric activity of the inhibitor and may shed light on how inhibition is transduced from the drug binding site through the pseudokinase domain.

We next identified resistance variants for a second inhibitor (NDI-034858) which binds the same allosteric pocket of TYK2 but in a different orientation and assessed whether we could use the spectrum of resistance variants identified by DMS to distinguish subtle differences in binding interactions between the two small molecules. Comparing drug-resistance profiles of matched (IC_99_) concentrations for BMS-986202 with NDI-034858 identified drug-resistant variants that are common to the two drugs, along with variants that uniquely ablate activity of one or the other (**Fig. 3E-G**, **Supplementary Fig. 5C**). For example, at position 671, which forms the pocket unique to TYK2 that is thought to drive target specificity of BMS-986202^54^, 11 variants at this position confer resistance to BMS-986202 (**Fig. 3F**). Only half of these variants also disrupt the activity of NDI-034858 (**Fig. 3F**). Conversely, variants at positions 603 and 694 disproportionately lead to resistance to NDI-034858 (**Fig. 3F**), and these sites have been shown to provide key interactions between TYK2 and this compound^46^. This analysis also highlights residues that are important for protein-drug interactions (e.g., residues 601, 640; **Fig. 3F**) that have not previously been shown to participate in direct interactions with either compound. Collectively, these results demonstrate that DMS comprehensively identifies variants that lead to drug resistance and provides an important layer of functional insight into protein-drug interactions that complements static structural data.

Lastly, we performed IFN-α signaling DMS assays at lower concentrations (IC_75_) of both inhibitors to identify variants that potentiate, or improve, the effect of the compounds (see **Methods** for details). In total, we identified 35 variants that increased inhibitor activity, including 10 that uniquely potentiate BMS-986202, 13 that uniquely potentiate NDI-034858, and 6 that potentiate both compounds (**Fig. 3H-J, Supplementary Fig. 5D**). While there was some positional overlap between variants that led to resistance or potentiation (e.g., compare position 758 results for NDI-034858 in **Fig. 3** panels **F** and **I**), the distributions were largely unique (**Fig. 3F-G, I-J**). Interestingly, there are three positions (residues 603, 694, and 739) where drug-potentiating variants of one drug occur at a position where there exists a known interaction with the other drug (**Fig. 3I, red boxes**). For example, N739 forms a known interaction with BMS-986202, and mutation of this residue to H or M increases the activity of NDI-034858. The converse is true of positions 603 and 694. By capturing potentiation at positions where a unique protein-drug interaction exists for just one compound, DMS reveals that there is capacity to recreate similar interactions with other small molecules. Traditional small molecule drug design employs structure-activity relationships (SAR) of small molecules, testing how iterative changes to a small molecule alter the function of a target. Our data captures the functional consequence of making changes to the protein instead, performing inverse SAR to highlight key residues on the target to exploit for drug design. Findings like these suggest that DMS data can be used to identify novel protein-drug interactions that could be reverse engineered into small molecules to increase compound potency, potentially saving time by reducing the number of iterations in the design process.

### Variants that protect against autoimmune disease reduce TYK2 abundance

Our TYK2 DMS experiments measure the effects of all common and rare human missense and nonsense variants on TYK2 signaling and abundance. This allows us to comprehensively assess how changes in TYK2 function relate to susceptibility to immune phenotypes, which could point to specific therapeutic modalities to prioritize in future TYK2 drug discovery programs. To this end, we examined how altering TYK2’s IFN-α signaling function or protein abundance relates to risk of autoimmunity.

We first focused on P1104A (rs34536443), the most well characterized of the common TYK2 variants, which has been shown to be protective broadly against autoimmune phenotypes^5,8,55^. Using genotype-phenotype information for a cohort of more than one million total individuals (from UK Biobank, FinnGen, and other sources), we performed a Phenome-wide association study (PheWAS) and independently confirmed that this allele is protective against many autoimmune diseases (**Fig. 4A**, **Supplementary Fig. 6A**; see **Methods** and **Table S2** for full list of data sources). While P1104A has been reproducibly shown to reduce TYK2 signaling through the interleukin-23 (IL-23) pathway^8,9^, its effect on IFN-α signaling has been inconsistent in the literature. Some studies report that it reduces IFN-α signaling activity^8^ and others report no effect^9^. P1104A showed no significant effect on TYK2 activity in our IFN-α assay (FDR > 0.01). This does not preclude there being an effect of the variant on IFN-α signaling in other conditions not tested here, but it does support hypotheses that P1104A has a lower impact on interferon than interleukin signaling pathways^9^. Although it showed no effect on interferon signaling, P1104A did, interestingly, result in significantly lower TYK2 protein abundance in our DMS assay (**Fig. 4B**; FDR < 2.2E-16).

**Fig. 4:**
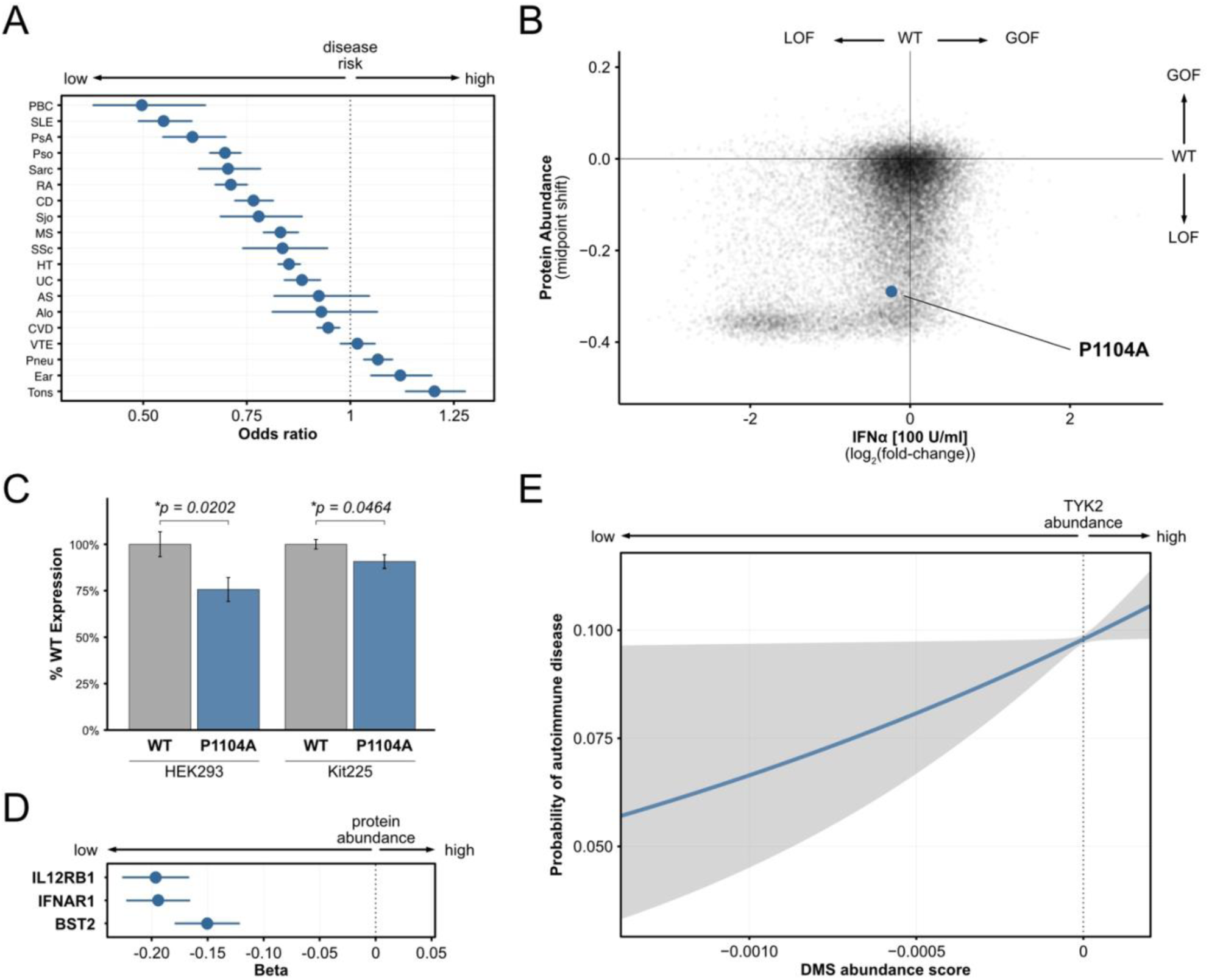
Human variants that protect against autoimmune disease reduce TYK2 abundance. **A**, Odds ratios from comprehensive PheWAS for the TYK2 P1104A allele (see **Methods**), which confirm protective effect on many autoimmune phenotypes (see **Supplementary Table S2** for list of phenotype abbreviations). **B**, Deep mutational scanning results for the IFN-α (x-axis) and protein abundance (y-axis) assays, as in Fig. 2B. Each point represents a unique missense or nonsense variant, and P1104A is highlighted in blue. GOF, gain-of-function; LOF, loss-of-function; WT, wild-type. **C**, TYK2 protein abundance measured using Western blotting of HEK293T and Kit225 cells harboring the wild-type (WT, gray) and P1104A (blue) alleles of TYK2. Data are shown normalized by the abundance of the wild-type allele for each cell type. Bars represent means, with error bars reporting standard error and *P* values determined by unpaired t-test. **D**, Effect sizes (*beta*) of association between P1104A allele and abundance of three proteins in human plasma from UK Biobank. Error bars are 95% confidence intervals. **E**, Rare variant dose-response curve for TYK2 variants found in the UK Biobank shows that TYK2 protein abundance (x-axis, from DMS assay) predicts the probability of autoimmune disease diagnosis (y-axis). See **Methods** for details. Gray shading indicates 95% confidence intervals.

The protein abundance DMS assay reports a relative, rather than absolute, effect on abundance and uses an engineered reporter. Therefore, we sought to validate this finding in several ways. First, we individually cloned and tested P1104A and a panel of other TYK2 alleles using the protein abundance reporter used for the DMS (see **Methods**). Consistent with the DMS library results, fluorescence activated cell sorting of the individual P1104A cell line validated that the allele led to lower normalized reporter signal than wild-type (**Supplementary Fig. 6B**; mean GFP:BFP signal for P1104A: 0.92; for WT: 1.16; p < 0.0001 by Welch’s t-test, n = 435,000 and 353,342 cells, respectively). Next, we integrated wild-type and P1104A TYK2, with no additional protein tags, under the control of a constitutive promoter in single copy at the H11 landing pad locus of HEK293T cells (see **Methods**). Western blot analyses of these lines showed that P1104A resulted in a ∼25% reduction in TYK2 abundance relative to wild-type (**Fig. 4C**, p = 0.0202 by unpaired t-test). Third, we used CRISPR/Cas9 engineering to generate a Kit225 cell line harboring the P1104A allele in homozygosity at the endogenous *TYK2* locus (**Supplementary Fig. 6C,** see **Methods**). Western blotting showed that the P1104A allele resulted in ∼10% lower TYK2 protein abundance than wild-type in Kit225 cells (**Fig. 4C, Supplementary Fig. 6D**, p = 0.0464 by unpaired t-test). Finally, we used a protein quantitative trait loci (pQTL) map within the UK Biobank^56^ to explore if P1104A (rs34536443) is associated with reduced abundance of any proteins in human plasma (see **Methods**). The abundance of TYK2 itself was not directly measured in this dataset, but P1104A is significantly associated with the reduced abundance of several proteins that were measured, including the interleukin receptor IL12RB1 and the interferon receptor IFNAR1 (**Fig. 4D, Supplementary Fig. 6E**, p < 5e-08, genome-wide significance via pQTL mapping as reported by UKB-Pharma Proteomics Project). These proteins are the two most significant pQTL associations for P1104A and are the receptors that directly interface with TYK2 in the IL-12/IL-23 and IFN-α signaling pathways, respectively^3^. Loss of TYK2 is well-established to lead to reduced localization of both receptors at the cell surface^2^. Collectively, these results support that the P1104A allele, which is protective for autoimmunity, leads to a modest but significant reduction of TYK2 abundance in both engineered cell lines and in humans.

A second common missense variant in TYK2 that changes the isoleucine at position 684 to serine (I684S; rs12720356) is also associated with a variety of autoimmune phenotypes. Its association is more complex than P1104A, providing protection against some phenotypes and risk to others^8^, which we confirmed through the independent PheWAS validation described above (**Supplementary Fig. 6A**). Like P1104A, I684S has no significant effect on IFN-α signaling in our DMS assay (FDR > 0.01) but does negatively impact protein abundance (FDR < 2.2E-16) (**Supplementary Fig. 6F**). This variant is also a significant pQTL for abundance of IL12RB1 and IFNAR1 in human serum (**Supplementary Fig. 6E**, p < 5e-08). Similarly to P1104A, these results support that common missense variants in TYK2 that are associated with autoimmune phenotypes lead to reduced TYK2 abundance.

Finally, we assessed if rare human *TYK2* variants that decrease IFN-α signaling or protein abundance show analogous protection against autoimmune phenotypes. Using autoimmune phenotype and *TYK2* genotype information from the UK Biobank, along with the DMS data described above, we constructed genetic dose-response curves^57^ comparing TYK2 function (either IFN-α signaling or protein abundance) with probability of having an autoimmune phenotype (see **Methods** for details). This uncovered a quantitative dose-related relationship between the probability of autoimmune disease and TYK2 abundance, but not IFN-α signaling function (**Fig. 4E**, **Supplementary Fig. 7**). In other words, reduced TYK2 abundance provides subtle but significant protection against developing autoimmune disease (**Fig. 4E**, p = 0.006, **Table S3**). Collectively, these results support that both common and rare sequence variants that reduce the abundance of TYK2 protein confer protection against autoimmune phenotypes. This strongly suggests that therapies, such as targeted protein degraders, that specifically reduce the abundance of TYK2 would be effective for the treatment of autoimmune diseases.

## Discussion

Over the last decade, deep mutational scanning has become a powerful technology for elucidating protein structure-function relationships and human variant interpretation. As such, it has substantial potential to improve many stages of drug discovery and development. However, this potential has yet to be fully realized because it requires highly quantitative methods that are directly relevant to the mechanisms underlying specific human phenotypes. In previous work, we described the development of quantitative DMS assays and analysis methods for GPCR proteins (β_2_AR and MC4R) and showed how they can elucidate structure-function relationships for the targets and their interactions with both native and novel agonists^22,23^. In this study we meaningfully extend these methods to another important target class for drug discovery (kinases) and uncover key insights into the biology of TYK2, a critical new target for the treatment of autoimmune disease.

First, we show how integrating DMS assays for multiple distinct functions (cytokine signaling and protein stability) can distinguish sites on TYK2 with important functional activities, such as allosteric regulation and protein-protein interaction. The ability to systematically map functionally critical protein regions and residues can influence early stages of drug discovery, where these data can be combined with structural insights to define relevant surfaces on the protein for drug targeting. Our DMS datasets successfully mapped the binding sites of small molecule allosteric inhibitors of TYK2 and distinguished subtle differences in protein-drug interactions at the amino acid level. These experiments capture the functional consequence of protein changes on drug activity, which serves as a valuable complement to traditional structure-activity relationship (SAR)-based methods where the protein is kept static and the small molecule design is iteratively changed. Such insights could be used to streamline compound optimization. Finally, we discovered that human *TYK2* variants that protect against autoimmune disease, including the common P1104A variant and many rare variants of previously unknown function, are associated with reduced TYK2 protein abundance. This finding strongly suggests that therapies designed to reduce TYK2 protein levels, such as targeted protein degraders, could be particularly effective for treating autoimmune diseases.

Collectively, this work demonstrates that high quality DMS data leads to actionable therapeutic insights that span multiple critical stages of drug discovery: defining a desirable pharmacological profile, identifying functionally relevant sites of action to target, and elucidating structure-activity relationships (SAR) to guide chemical optimization. The ability to inform drug discovery and development at multiple stages highlights the value of DMS as a tool for integrating structural, biochemical, and genetic information toward the development of better therapies. As DMS technology continues to improve, it will increasingly provide actionable insights for drug discovery and precision medicine.

## Materials and Methods

### Reporter design and optimization

The IFN-α signaling reporter was adapted from DMS assays used previously for GPCR signaling^22,23^ that employ RNA sequencing of transcriptional reporters to assess variant effects. Briefly, the plasmid design for this system was modified to include a doxycycline-inducible *TYK2* allele transgene upstream of an eight copy Interferon-Stimulated Response Element (ISRE)^24^, which was followed by the Promega minimal promoter (minP) and a *luciferase* reporter gene harboring a unique DNA sequence barcode in the 3’ untranslated region.

The protein abundance reporter was modified from the previously described VAMP-seq reporter system^25^. This system, which uses a polycistronic construct containing an amino-terminal GFP-tagged target protein (in this case TYK2) and a second fluorescent protein to normalize protein abundance to transcription levels, employs fluorescence-activated cell sorting (FACS) and DNA sequencing to assess variant effects. We made three primary design modifications to this system. First, the abundance reporter was integrated into a landing pad site engineered at the *H11* safe-harbor locus to include a CAG promoter^22^, rather than a doxycycline-inducible Tet promoter, leading to constitutive expression of the reporter. Second, we used a co-transcriptionally linked BFP instead of mCherry for normalization, as our base cell line already expressed mCherry. An Internal Ribosome Entry Site (IRES) allowed for independent translation of the BFP from the shared reporter transcript. Finally, the full-length reporter plasmid included a Blasticidin resistance gene (BSR) transcriptionally linked to the BFP by a self-cleaving T2A peptide (GFP-TYK2-IRES-BFP-T2A-BSR) so that Blasticidin selection, rather than fluorescence-activated cell sorting (FACS), could be used to remove cells without the reporter construct integrated correctly into safe-harbor locus.

### Base cell line generation

All copies of the *TYK2* gene were knocked out in HEK293T cells harboring a single copy Bxb1 recombination-based landing pad at the *H11* locus^23^ using a CRISPR multi-guide RNA deletion strategy with reagents from Synthego (Gene Knockout Kit v2). Single guide RNA (sgRNA) sequences were as follows: sgRNA1-5’-CCACUGUCCCGGAUGUAGCA-3’; sgRNA2: 5’-CACACGCUCUGUGCCGAAGC-3’; sgRNA3: 5’-UUUGACCAUGACCAUCUGCU-3’. Guide RNAs were mixed with Cas9 protein to form ribonucleoproteins (RNPs), which were then transfected into cells using Lipofectamine CRISPRMAX reagents (Invitrogen). Knockout efficiency was measured at both 7– and 19-days post-transfection (passages 1 and 6, respectively) by performing PCR with validation primers (FWD: 5’-AACACAGAAACCCACCCCAG-3’; REV1: 5’-CCCAGCTTCAAGGACTGCAT-3’; REV2: 5’-CTCCTTCCGCCGGCATATC-3’) on genomic DNA extracted from pools of transfected cells (Zymo Quick-DNA, #D3024). The presence of multiple laddered bands in treated cells confirmed successful editing at the *TYK2* locus. Sanger sequencing was also performed on the PCR products, and Synthego’s ICE analysis tool was used to confirm an editing efficiency of 80% or greater for both the positive control and pooled *TYK2* knockout transfections. Isoclonal cell lines were obtained by seeding the edited cell pool into ClonaCell (STEMCELL, #03814) in a 10-cm dish. Cells were grown for 20 days, and then single colonies were picked into a 96-well plate in Dulbecco’s Modified Eagle Medium (DMEM; Gibco) + 10% dialyzed fetal bovine serum (FBS; Gibco). Isoclonal cultures were grown for an additional 2-3 weeks, during which two rounds of genotyping were performed by Sanger sequencing to confirm gene editing (using the primers listed above and the Zymo ZR-96 PCR kit, #D4024). In sequence-validated isoclones, western blotting was used to validate that no TYK2 protein was produced, further confirming editing of all copies of *TYK2* (see **Western blotting** methods section, below, for detailed methods). Finally, isoclonal cell lines were tested by luciferase assays using transient delivery of the IFN-α reporter plasmid described above (see ***Reporter design and optimization***) and plasmids containing transgenes for *STAT2* and *IRF9*, which are required for IFN-α but not expressed in HEK293Tcells. The best knockout cell line was chosen for a combination of low background activity in transient reporter assays and healthy growth characteristics.

We then engineered IFN-α signaling activity into the *TYK2* knockout cell line by performing stable piggyBac transposon-based integration of separate plasmids constitutively expressing the *STAT2* and *IRF9* genes under the control of the EF1A promoter (NM_005419.4 and NM_006084.5, respectively). The resulting cells were then isocloned as described above. Clonal lines were tested for signaling assay efficiency by Bxb1-mediated integration of the IFN-α reporter constructs described above (see ***Reporter design and optimization***) into the *H11* landing pad locus. With this reporter system, stimulation of cells with IFN-α cytokine leads to TYK2-dependent phosphorylation of STAT1 and STAT2 proteins, followed by transcriptional activation of the ISRE element by the STAT1/2-IRF9 complex. The reconstitution line with the best signal to noise for TYK2-dependent activation (determined by calculating Cohen’s D and Z-factor scores of luciferase luminescence with and without doxycycline induction of TYK2) was chosen as the base cell line for all DMS libraries (**Supplementary Fig. S1A**).

### DMS library generation and barcode-mapping

DMS plasmid and cell libraries for both assay types were generated as previously described^22,23^. In brief, the *TYK2* coding sequence (P29597) was first divided into 17 segments (“sub-libraries”) of ∼210 base pairs (bp), and pools of ∼270 bp long DNA oligos (Twist Bioscience) were designed to encode every possible single amino acid substitution to the *TYK2* coding sequence. Each *TYK2* coding segment was amplified from the DNA oligo pools and cloned into base vectors through a multi-step process to yield pooled libraries of *TYK2* variants with fully intact and barcoded reporter gene cassettes (see **Table S4** for all amplification and sequencing primers used in this process). First, DNA barcodes were appended to variant alleles during the initial amplification step of each of the 17 oligo sub-libraries at sufficient scale to yield many unique barcodes per variant. These variant-barcode linked PCR amplicons were cloned into sub-library specific destination plasmids. After this first step of cloning, variant segments were amplified, along with the associated barcode, and sequenced with 2×150 paired-end reads on an Illumina NextSeq 2000 instrument using a 300-cycle P3 kit. Sequencing data was de-multiplexed and variant-barcode associations were mapped as previously described^23^. A second cloning step was performed to integrate the missing piece of the reporter plasmid between the variant allele and barcode, resulting in fully assembled plasmid libraries. These were then co-transfected with a plasmid encoding Bxb1 recombinase, into the HEK293T cell line described above, and properly targeted cells were selected with 10 µg/ml Blasticidin (Gibco) for ∼14 days. This resulted in a library of HEK293T cells, with each cell containing a single *TYK2* allele-barcode combination integrated in single copy in a landing pad at the *H11* safe harbor locus.

### DMS assay execution and sequencing

IFN-α signaling experiments were performed in triplicate as described for previous transcriptional DMS reporters^23^ with some modifications: DMS cell libraries were seeded at ∼19E6 cells per 150 mm tissue-culture treated dish for each assay replicate. TYK2 expression was induced 24 hours later by replacing media with 24 mL DMEM + 0.5% dialyzed FBS +/− 1 µM doxycycline (ApexBio). Signaling was induced 24 hours later by adding human Interferon-Alpha 2a cytokine (“IFN-α”, PBL Assay Science #11100, see **Table 1** for concentrations used for each experiment). For experiments involving TYK2 inhibitors (MedChemExpress), a pre-drugging phase was introduced after overnight doxycycline induction where media was replaced with DMEM + 0.5% dialyzed FBS, +/− drug for 1 hour, followed by an exchange into DMEM + 0.5% dialyzed FBS, +/− cytokine, +/− drug. Six hours after cytokine stimulation, cells were harvested by scraping in 4 mL lysis buffer (RLT buffer [QIAGEN] + 143 mM β-ME). Cells were lysed, RNA was extracted from 1 ml of the homogenized lysate, and reverse transcription was performed on the extracted RNA using SuperScript IV Reverse Transcriptase (Invitrogen, #18091050). cDNA from each sample was treated with 3.2 µL RNase A (100 µg/ml, Thermo Fisher) and 8 µL RNase H (5000 U/mL, NEB) at 37 °C for 30 min, followed by concentration to ∼50 µL by spinning through Amicon Ultra 10 kDa concentrators (EMD Millipore, #UFC5010) for ∼8 min.

The TYK2 protein abundance DMS assay was performed by adapting the VAMP-seq technique^25^, with modifications as follows. Before performing the assay, the DMS HEK293T cell sub-libraries were bottlenecked to reduce barcode complexity to ∼20 barcodes per variant by seeding ∼200,000 cells from each sub-library into a single well of a 12-well plate followed by expansion. At the beginning of the assay, the bottlenecked sub-libraries were pooled, and each assay replicate was seeded with enough cells to achieve at least 7E6 cells per sub-library. Three assay replicates were seeded and independently sorted via FACS on three different days. Prior to sorting, cultures were harvested in log-phase and resuspended in *cold* DMEM + 10% dialyzed FBS at a density of ∼10E6 cells/mL. Cell slurry was passed through a 35 µm snap-cap filter into a 5 mL tube (Corning, #352235) 1 mL at a time to remove cell clumps and debris and then kept on ice until flow cytometry. Flow sorting was performed on a Sony MA900 using a 100 µm sorting chip. Control HEK293T cell lines (background reporter, BFP-only, and GFP-only) were first used to establish appropriate gates for forward scatter (FSC; single cells and no debris), back scatter (BSC; live cells), and fluorescence dynamic range for both BFP and GFP. Then, ∼100,000 cells from the pooled DMS library were flowed, and the fluorescence distribution was used to generate polygonal gates that divided the population into equal quartiles based on the GFP:BFP fluorescence ratio. These quartiles were assigned to the four collection bins, and 5 mL collection tubes containing 2 mL of DMEM + 20% dialyzed FBS were placed in each bin. The complete library was sorted, regularly replacing collection tubes as needed and keeping sorted cells on ice, until ∼25E6 cells per bin were collected. Sorted cells for each quartile bin were pooled into a 50 mL conical, spun for 5 min at 300 x g, and resuspended in 1.2 mL PBS. DNA was extracted from cells with the DNeasy Blood and Tissue extraction kit (Qiagen, #69504) using one purification column for every 5E6 cells. After addition of Buffer AL and ethanol, a shearing step that involved passing cell slurry 5x through an 18G syringe needle was performed to increase lysis before proceeding with genomic DNA extraction.

Both the IFN-α and abundance DMS libraries were assayed in biological triplicate and prepared for sequencing as previously described with minor modifications^23^. In brief, qPCR reactions were performed on 1 µL DNA (diluted 1:8 in water to avoid late-cycle assay inhibition) with NEBNext Ultra II Q5 polymerase (NEB, # M0544L) and SYBR Green (Thermo Fisher, #S7563) (see **Table S4** for qPCR amplification and sequencing primers described in this section). Illumina sequencing libraries were amplified with the same polymerase under the following cycling conditions: 98 °C for 30 s, X cycles of [98 °C for 10 s, 65 °C for 75 s], followed by an extension of 65°C for 5 min. The number of amplification cycles for each sample was set to the respective C_q_ value determined by qPCR. Amplified products were purified with SPRI beads (Beckman Coulter, #B23317). An additional round of qPCR and library amplification was performed with secondary primers to append sequencing indices and further amplify the sample libraries. This secondary amplification was performed in duplicate for each biological replicate to account for potential PCR amplification or sequencing errors. To account for differences in library yields, 3 µL of each amplified DNA library sample was run on a 4% E-Gel (Thermo Fisher) and densitometry was performed with Fiji^58^. Samples (i.e., from different replicates and assay conditions) were mixed at equimolar ratios into a single pool and then thoroughly purified by performing a SPRI-bead cleanup (1:1 ratio of beads:DNA, two 70% ethanol washes, and elution into 50 µL IDTE pH 8.0), agarose gel electrophoresis and gel extraction (elution into 36 µL water), and then another SPRI-bead cleanup (3:1 ratio of beads:DNA, two 70% ethanol washes, and elution into 36 µL IDTE pH 8.0). The purified library was quantified with the DeNovix High-Sensitivity Fluorescence kit and prepared for sequencing with a 30% PhiX spike-in. The final library mixture was sequenced using custom read and index primers (**Table S4**) on an Illumina NextSeq 2000 with a 50-cycle P3 kit.

### DMS primary variant effect analysis

Primary variant effect analysis of IFN-α signaling DMS experiments was performed as previously described^23^. Briefly, the first 21 bp of each single-end read was extracted and joined to the previously generated variant-barcode map (see ***DMS library generation and barcode-mapping***), requiring a perfect barcode sequence match. The combined mapped counts across all samples were analyzed using the same negative binomial generalized linear mixed model as the previous work, fitting one model per position and incorporating all variant counts at that position, as well as all wild-type counts from the corresponding protein segment. Using this model, we extracted log_2_(fold change) estimates for all variants (relative to wild-type TYK2) and associated standard errors. Then, for each assay, we computed the linear contrast of each variant in each condition against that same variant in the Untreated (no cytokine stimulation) condition, to obtain final variant effect estimates.

For abundance DMS analysis, we developed a new extension of our previous model that leveraged the multi-barcode library design to quantitatively capture the effect of variants on the FACS distribution of cells. We applied the same model as previously described^23^, except using the four fluorescence bins (25/50/75/100) as the “condition” and removing the offset. Then, for each variant, we computed a weighted average and standard error by simulating 1000 draws from a normal distribution, with mean and standard deviation defined by the estimate and standard error from the initial regression, respectively. This allowed us to compute a weighted average across all bins, as well as the corresponding standard error. The weights for each bin were specified as the mid-point of that bin (0.125/0.375/0.625/0.875). The resulting “midpoint” estimates and standard errors were used for downstream analysis.

### Orthogonal variant classification data

Variant effects for all reported TYK2 alleles available were downloaded from ClinVar^38,59^ on 09 January 2025.

### Classification of signaling-only LOF variants

To define TYK2 variants that affect signaling but not abundance, the 100 U/mL IFN-α signaling and protein abundance DMS datasets were merged and then filtered to identify variants that met both of the following criteria:

1. Significant signaling LOF: log_2_(fold change of variant relative to wild-type) in IFN-α condition < 0 *<u>and</u>* FDR(IFN-α condition) < 0.05
2. No statistically significant negative effect on Abundance: Abundance midpoint shift > 0 *<u>or</u>* FDR(midpoint shift) > 0.1

### Identification of drug resistance and potentiation variants

To define TYK2 variants that conferred drug resistance or potentiation, drug-treated conditions (100 U/mL IFN-α stimulation + inhibitor, see **Table 1**) were assessed for variants that led to a significantly different drug-response relative to WT (FDR < 0.01). For example, to define variants resistant to BMS-986202 inhibitor, we filtered for variants that were significantly *more active* than WT, indicating resistance to inhibition (**Fig. 3B—D**). To define drug-potentiating variants, we filtered for the opposite: variants that had significantly *less activity* than WT at sub-maximal inhibitor dose (IC_75_), indicating a stronger response to inhibitor. For potentiation profiling drug-treatment conditions were normalized to the cytokine stimulated condition without drug treatment to focus on drug-specific effects.

### Data and code availability

Raw FASTQ files for each DMS assay sample are available on SRA in the BioProject with accession PRJNA1291213. Complete oligonucleotide-barcode maps for the DMS libraries, and mapped barcode-level read counts for each assay sample set, are available as tab-delimited text files on Zenodo at accession 15347448.

Code for data processing, statistical modeling, and summary statistics are available on GitHub at https://github.com/octantbio/bms-dms. We provide one summary statistics file per dataset, with all variant effects in each condition, and a separate single file covering all assays containing the linear contrasts of each variant effect in each condition against the same variant in the Untreated control. For the abundance dataset, we provide the raw variant effect summary statistics, as well as the post-processed “midpoint” statistics.”

### Structural modeling

Molecular visualization of variant effects on TYK2 was performed with UCSF ChimeraX^60^. For **Figs. 1C** and **2D**, the cryo-EM structure of the TYK2 Kinase-Pseudokinase domains (PDB: 4OLI) was used, and blue amino acid positions represent those with significant LOF variants. The full-length composite structure of TYK2 (**Fig. 2E**) was generated from structures of TYK2(FERM-SH2)-IFNAR1 (PDB: 4PO6) and TYK2(Kinase-Pseudokinase) (PDB: 4OLI), using full-length JAK1-IFNAR1 (PDB: 7T6F) as a scaffold for the ChimeraX MatchMaker tool. For visualizing variant effects on protein-ligand interactions (**Fig. 3 and Supplementary Fig. 5**), crystal structures of the inhibitor-bound pseudokinase domain were used (PDB: 6NZP and PDB: 8S9A), and residues were colored pink, cyan, or orange to denote association with BMS-986202, NDI-034858, or both, respectively.

## GWAS and fine-mapping *TYK2* signals

We analyzed the association of P1104A with autoimmune diseases, utilizing GWAS data aggregated together across a variety of sources, including the UK BioBank, FinnGen, and large studies reported in the literature (**Table S2** lists all data sources for each phenotype)^61–69^. When multiple, non-overlapping studies were available, we performed inverse-variance fixed-effects meta-analyses to obtain the most robust effect size estimates for P1104A on the diseases. We selected autoimmune diseases for inclusion in our analysis based on the availability of well-powered GWAS. We additionally included cardiovascular diseases and several infectious diseases that could be utilized for safety considerations.

We conducted association testing with data from the UK Biobank to validate previous findings that common genetic variants in *TYK2* are associated with autoimmune diseases^8^. We initially selected variants for inclusion in our analysis, selecting high-quality common variants (imputation quality INFO score > 0.8, minor allele frequency > 0.01) at the *TYK2* locus (including the 200 kb region surrounding *TYK2*). Using these variants, we conducted association analyses with the autoimmune traits in UK Biobank that have enough cases to perform a well-powered analysis, along with the pooled autoimmune trait. We used UK Biobank for this purpose due to the ability to utilize individual level data. The individual traits considered were Psoriasis (PsO), Rheumatoid Arthritis (RA), Hypothyroidism (HT), Inflammatory Bowel Disease (IBD), and Ulcerative Colitis (UC). Covariates were the same as in our rare-variant analysis (first 10 principal components, sex, age). REGENIE was used for all association testing^70^.

Downstream of the association testing, we first dissected independent signals at the locus with conditional analysis, using the GCTA-COJO software^71^. Firstly, we used the 5 x 10^−8^ p-value threshold to obtain independent conditional signals in the investigated region. We then used the fine-mapping approach SuSIE^72^ to confirm the most likely causal variants at these loci using probabilistic annotations. Furthermore, we conducted conditional analysis using a lowered threshold (p-values < 5 x 10^−5^) to capture additional ‘suggestive’ signals and to shed light on potential signals that we were underpowered to detect.

Consistent with previous analyses^8^, P1104A (rs34536443) was the most likely causal variant for the primary signal for Psoriasis, RA, HT, and the pooled autoimmune trait (**Supplementary Fig. 6A**). It was also the most likely causal variant for a secondary suggestive signal for UC and IBD. Meanwhile, another missense variant, I684S (rs12720356), was identified as the most likely causal variant for the primary signal for IBD and UC, and the secondary signal for Psoriasis. The directionality for I684S is opposite for Psoriasis (protective) versus IBD/UC (risk), also consistent with previous reports^8^. In addition to these, we see a secondary suggestive intronic signal for hypothyroidism, with rs11085727 prioritized as the most likely causal variant here. It should also be noted that, while I684S does not come up as suggestively significant for RA, it barely misses the threshold (conditional p-value: 8 x 10^−5^).

### UK Biobank genetic data

To build genetic dose response curves^57^ with IFN-α and TYK2 protein abundance DMS data, we first identified rare human *TYK2* missense or nonsense variants in the UK Biobank 450k whole exome sequencing dataset^73^. We annotated variants using the Ensembl Variant Effect Predictor^74^ and filtered to canonical transcript missense variants. We identified 447 rare protein-altering variants (MAF < 0.01%) and used these variants to both calculate per individual DMS scores and perform rare variant analyses testing for an association of *TYK2* with autoimmune disease.

### UK Biobank phenotype data

We first filtered UK Biobank participants by excluding participants with outlier heterozygosity or missing rates, participants where submitted sex did not match genetically inferred sex, and participants with putative sex chromosome aneuploidy. Next, we subset to participants with British White ancestry and included only one individual per related pair. We then identified participants with Psoriasis, Rheumatoid arthritis, Systemic lupus erythematosus, Ankylosing spondylitis, Primary biliary cirrhosis, Ulcerative colitis, and Crohn’s disease by querying participants’ diagnosis data with a combination of ICD10 codes, general practice records, and PheCode definitions (**Table S5**). We pooled the participants with any endpoint diagnosis into a general *TYK2* associated autoimmune disease cohort of 31,726 total cases to maximize association testing power. To generate an “unaffected” control cohort, we started with the general quality control filtered set of participants in UK Biobank and excluded all participants with the autoimmune diagnoses listed above. This resulted in a control cohort of 308,674 participants.

### Human variant analysis using DMS scores

We used the DMS IFN-α and protein abundance data to calculate a per individual DMS score, analogous to a polygenic risk score where the variant effect size is its DMS scaled log_2_(fold change relative to wild-type) value. We calculated DMS scores for each participant (j) given their genotype (G) at each *TYK2* variant (i) as,

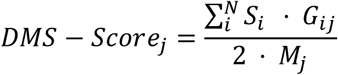

where S denotes the scaled log_2_(fold change relative to wild-type) score for a variant and M denotes the total number of non-missing genotypes for TYK2 variants.

We aimed to test if DMS scores for variants that are rare in the human population could predict general autoimmune disease case status, and so we fit logistic regression models using the IFN-α and protein abundance DMS scores as predictors of disease case status. The common deleterious *TYK2* missense variant rs34536443 (P1104A) is strongly associated with protection from various autoimmune diseases^8^, and we anticipated that the effect of common variants in linkage disequilibrium with the rare variants used to construct the DMS score could confound the analysis. We used a conservative minor allele frequency filter (MAF < 10^−4^) to control for these common variant haplotype effects, but we also explicitly included participant rs34536443 allele dosage as a covariate in the model in addition to including PCs 1-10, sex, and age. We fit a series of logistic regression models with increasing complexity, as well as a null model including only covariates, and used Akaike information criterion (AIC) to select the best fit model (see **Tables S3, S6-S8** for model selection details and coefficients).

### Rare variant association test

As a comparison to our novel DMS score association testing, we also employed traditional rare variant association analyses. We tested for associations using two methods that each assume different patterns of variant effects on the disease phenotype: a variance component test and allelic series test. We implemented the variance component tests with the R SKAT-O package^75^, and the allelic series test with the R AllelicSeries package^76^. We included PCs 1-10, sex, and age as covariates in all rare variant association tests. We implemented all three association test approaches for each disease endpoint individually and for the general autoimmune disease super set.

The allelic series test requires variants to be categorized based on their predicted effect, as either benign missense variants, deleterious missense variants, or protein truncating variants. We classified “missense variants” in the allelic series test if the Variant Effect Predictor (VEP) consequence was missense variant, in frame deletion, in frame insertion, stop lost, start lost, or protein altering. We classified protein truncating variants as stop gained, splice acceptor variants, splice donor variants, or frameshift variants. To further subset missense variants, we classified variants as “benign” if predicted to be “tolerated” by SIFT or if they have a CADD Phred score less than or equal to 20. We classified variants as “deleterious” if predicted to be “damaging” by SIFT or if they have a CADD Phred score greater than or equal to 20. We further filtered protein truncating variants to only include LOFTEE High Confidence predicted loss-of-function.

### Cell lines harboring individual alleles of TYK2 for variant effect validation

To confirm the effects of TYK2 variants on IFN-α signaling and protein abundance, individual variants were cloned and integrated into the *H11* landing pad of our HEK293T reporter cell line following the same methods as DMS library integrations described above. N-terminally GFP-tagged *TYK2* constructs were used to validate protein abundance effects by flow cytometry (**Supplementary Fig. 6B**). Untagged *TYK2* constructs were used to validate protein abundance effects by western blotting. (**Supplementary Fig. 6D, Supplementary Fig. 8**). Both tagged and untagged constructs had the same architecture as the abundance DMS reporter described above, with *TYK2* expression driven by a CAG promoter.

### FACS-based validation of protein abundance

Cell lines harboring a single copy of GFP-tagged TYK2 variants were grown to log-phase, harvested, and resuspended in *cold* DMEM + 10% dialyzed FBS. Cell slurry was passed through a 35 µm snap-cap filter into a 5 mL tube (Corning, #352235) to remove cell clumps and debris and then kept on ice until flow cytometry. Cells were processed in the same manner as the protein abundance DMS experiment, collecting BFP and GFP signal per cell (but without subsequent cell collection). Approximately 600,000 events were collected for each sample, and FloJo (BD Biosciences) was used to generate histograms of the GFP:BFP ratio of each sample (**Supplementary Fig. 6B**).

### Engineering the P1104A allele into endogenous *TYK2* of Kit225 cells

CRISPR RNP complexes containing an sgRNA with targeting sequence CTCCAGCCAGAGCCCCCCCA and *S. pyogenes* HiFi Cas9 protein V3 (IDT) were delivered to cells, along with an ssODN homologous-driven repair (HDR) donor (GGTATGCCCCAGAGTGCCTGAAGGAGTATAAGTTCTACTATGCGTCAGATGTCTGGTCCTTCGGGGTGACCCTGTATGAGCTGCTGACGCACTGTGACTCCAGCCAGAGCGCCCCCACGGTGAGAGCCAGGCCCGCAGCCCCACCGGGAG) (IDT) with “AltR” end-modifications, using Nucleofection (Lonza) in solution SF with pulse code CA-137. Following several days out-growth, the cells were single cell cloned by limiting dilution in 96 well plates. The resulting clones were replicated and screened for knock-in by amplicon next-generation sequencing (NGS). Briefly, PCR primers TYK2-P1104-2F (5’-GCCTAGCCAATGAGCATATCAC-3’) and TYK2-P1104-2R (5’-TCTCTCTCCGTCCTGTCCTGTCTTA-3’) were used to amplify the targeted region out of genomic DNA lysates, the amplicons were purified with AMPure XP beads, and sequencing libraries prepared using Illumina DNA Prep with Tagmentation. NGS sequencing libraries were pooled and sequenced on a MiSeq (Illumina) using 150 pair-end reads with dual indexes. Data was demultiplexed using bcl2Fastq and run through the CRISPR-DAV analysis pipeline as previously described^77^ with the HDR_NewBase parameter set to 10352442C (for the Chr19 change in the hg38 human reference genome) to specifically identify clones homozygous for the P1104A mutation. Kit225 cells were grown in RPMI 1640 (containing L-glutamine) with 10% FBS, 20 ng/mL human recombinant IL-2 (Thermo Fisher, #PHC0023).

### Western blotting

For western blot experiments, cells in log-phase growth were harvested by lifting and washing 3x with cold PBS before freezing dry cell pellets at –80° C. Cell pellets were resuspended with 100 µL Lysis Buffer (RIPA, Thermo Scientific #89901; 1 µL Universal Nuclease, Pierce #88701; 1x Halt Protease and Phosphatase Inhibitor Cocktail, Thermo Scientific #78446) per 5E6 cells and lysed by incubation for 60 min on ice, vortexing every 10 min. Cell lysates were clarified by spinning for 10 min at 13,000 rpm and 4° C, and the resulting supernatant was quantified using a Rapid Gold BCA Protein Assay Kit (Pierce, #A53225). Lysate was diluted to between 1 and 20 μg (depending on the experiment), and 5 μl of lysate sample was mixed with LDS Sample Buffer (Invitrogen, #B0007) and Reducing Agent (Invitrogen, #B0009) before heating at 95° C for 5 min. Denatured samples were loaded into 4-12% Bis-Tris Plus mini gels (Invitrogen, #NW04125BOX) and run at 120 V for 70 min in 1x MOPS Running Buffer (Invitrogen, #B0001). Protein was transferred to a PVDF membrane (pre-activated with methanol for 5 min) with pre-chilled 1x Running Buffer (Invitrogen, #BT0006) using a Mini Blot Module (Invitrogen, #B1000) run at 20V for 70 min. Membranes were rinsed 3x with TBST (LICOR, #927-65001) before blocking overnight in Blocking Buffer (LICOR, #927-60001). The blocked membrane was cut just above loading control molecular weight (MW) band and incubated with primary antibodies diluted in TBST (1:500 rabbit anti-TYK2 [Abcam, #ab223733]); 1:10,000 mouse anti-GAPDH [Novus, #NB300-324]) for 1 hour at 25 °C on a rocker. Membranes were washed 3x with TBST before incubating with 1:15,000 dilutions of anti-rabbit or anti-mouse secondary antibody (LICOR, #926-32211 and #926-32210) in TBST. Final washes with 3x TBST and 3x TBS were performed before imaging with an Odyssey system (LICOR, #9120). Imaged blots were analyzed with Fiji^58^. For each lane, the integrated density of the full-length TYK2 band was normalized to the integrated density of the GAPDH loading control. The average GAPDH-normalized value for the WT allele was set to 100% for each blot, and this average was used to compute the GAPDH-normalized relative expression of all TYK2 bands in the same blot (both WT and P1104A samples). Lanes were excluded from analysis if there were clear blemishes near bands of interest, no signal was detected for either band, or either TYK2 or GAPDH were outside of the linear range of detection (**Supplementary Fig. 8**).

### Plasma pQTL association testing

We used data from the UK Biobank – Pharma Proteomics Project (UKB–PPP)^56^ to detect proteins whose plasma levels are associated with variants predicted to be loss–of–function for TYK2. To this end, we utilized the full pQTL mapping summary statistics from the UKB-PPP Consortium. Using the summary statistics for all 2,936 proteins from the UKB-PPP Olink panel, we analyzed whether P1104A and/or I684S was implicated as a causal variant for each protein’s plasma levels.

We utilize a similar procedure as with the GWAS (see **GWAS and fine-mapping *TYK2* signals**) to investigate causal variants, initially conducting conditional analysis for each protein to dissect conditionally independent signals at loci. The significance threshold for this analysis was at the 5E-08 level (genome-wide significance). Next, we used SuSIE^72^ to confirm the prioritized variants corresponding to these signals.

### Cell line validation

STR profiling performed by the University of California Berkeley DNA Sequencing Facility confirmed that cell lines used for DMS were HEK293-derived. Cell lines were verified as negative for mycoplasma contamination using the EZ-PCR Mycoplasma Detection Kit (Sartorius, #20-700-20).

### Figure generation

The following figures were generated using BioRender: Fig. 1E (full schematics), Fig. 2E (the lipid layer image only; the protein structure was generated using ChimeraX as described above).

## Acknowledgements

Molecular graphics and analyses performed with UCSF ChimeraX, developed by the Resource for Biocomputing, Visualization, and Informatics at the University of California, San Francisco.

## Ethics Declaration

This work was funded by Bristol Myers Squibb.

Competing Interests: CJH, NSA, RRW-T, CR, EMT, MM, BLJ, SK, and DED are current employees and/or share or options holders of Octant, Inc. ERH, KC, AH, GAM, AC, CN, KNW, SCW, JCM, PRS, and RMP are current employees and/or shareholders of Bristol Myers Squibb. All other authors have no competing financial interests.

## Supplementary Figures

**Supplementary Fig. 1:**
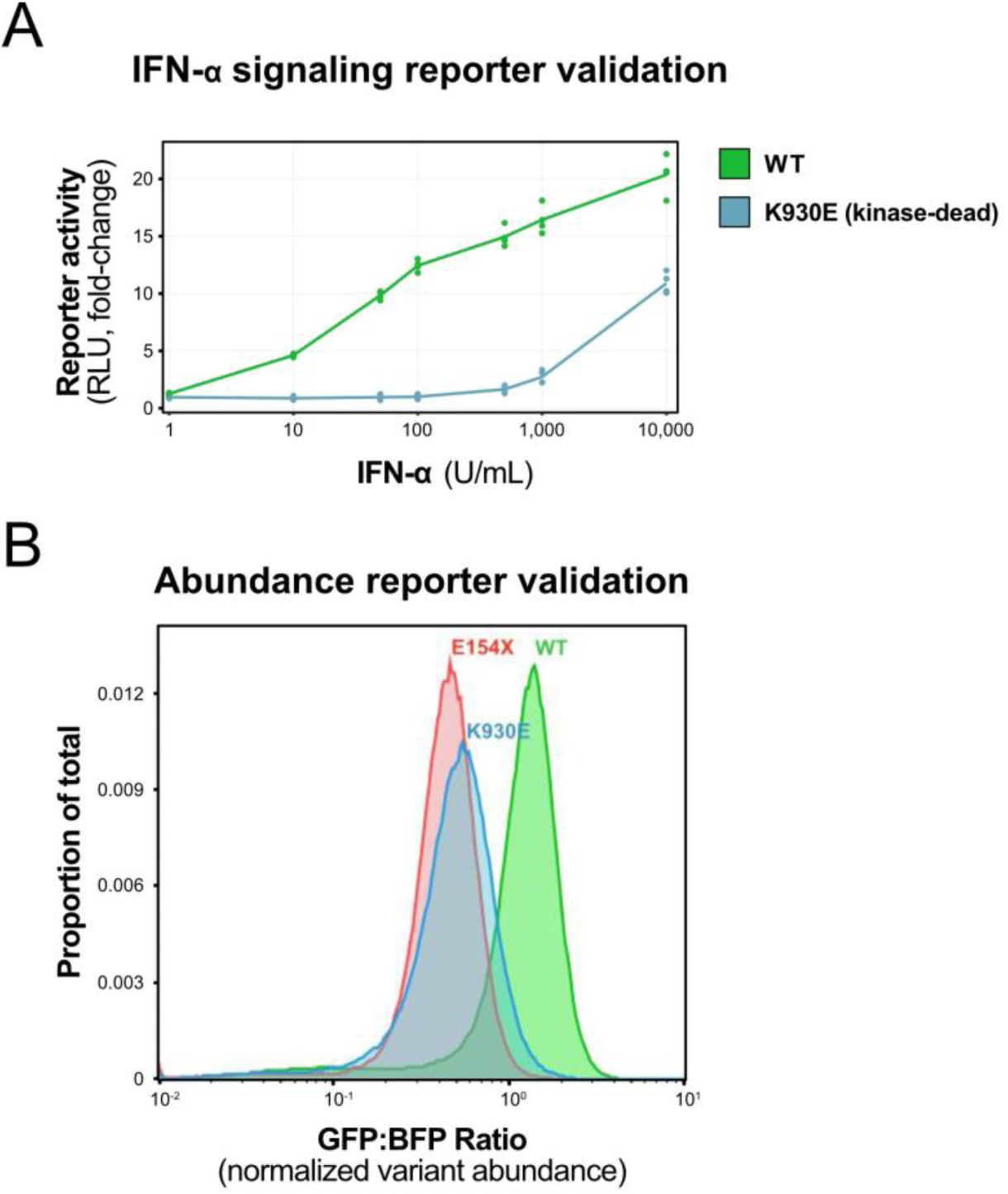
IFN-α and abundance DMS reporters are TYK2-dependent. **A**, IFN-α reporter assay dose-response curves for wild-type (WT) and catalytically inactive (K930E variant) TYK2. RLU, relative luciferase units. **B**, Flow cytometry data showing that the abundance DMS reporter distinguishes between WT and two strongly destabilizing variants (E154X and K930E).

**Supplementary Fig. 2:**
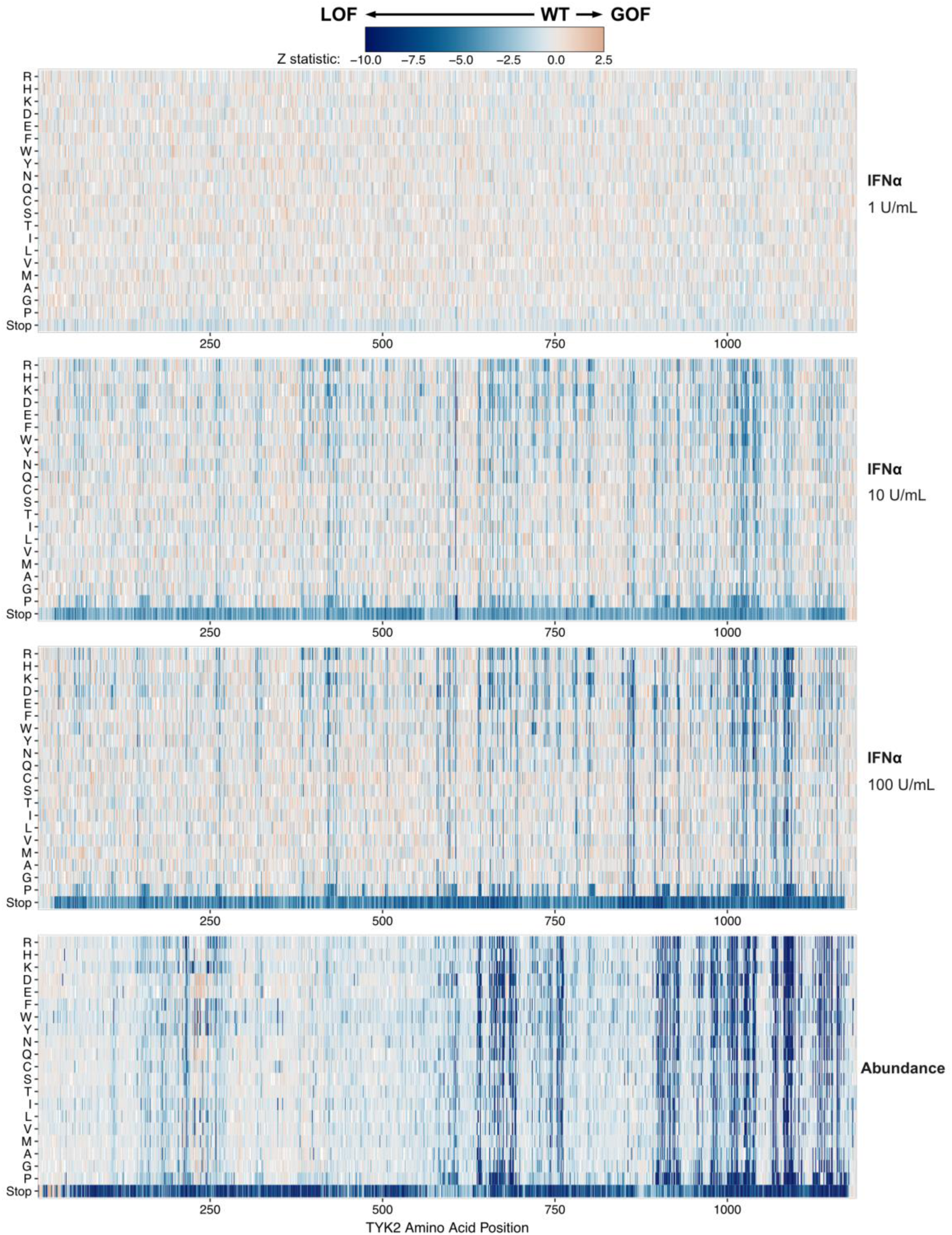
Results of IFN-α and abundance DMS assays. Heatmap showing results of the IFN-α (1, 10, and 100 U/mL stimulation conditions) and abundance DMS datasets generated in this study. Each cell represents a single variant allele colored by the magnitude of variant effect (Z-statistic). For each heatmap, the x-axis represents amino acid position, and the y-axis represents the induced amino acid change.

**Supplementary Fig. 3:**
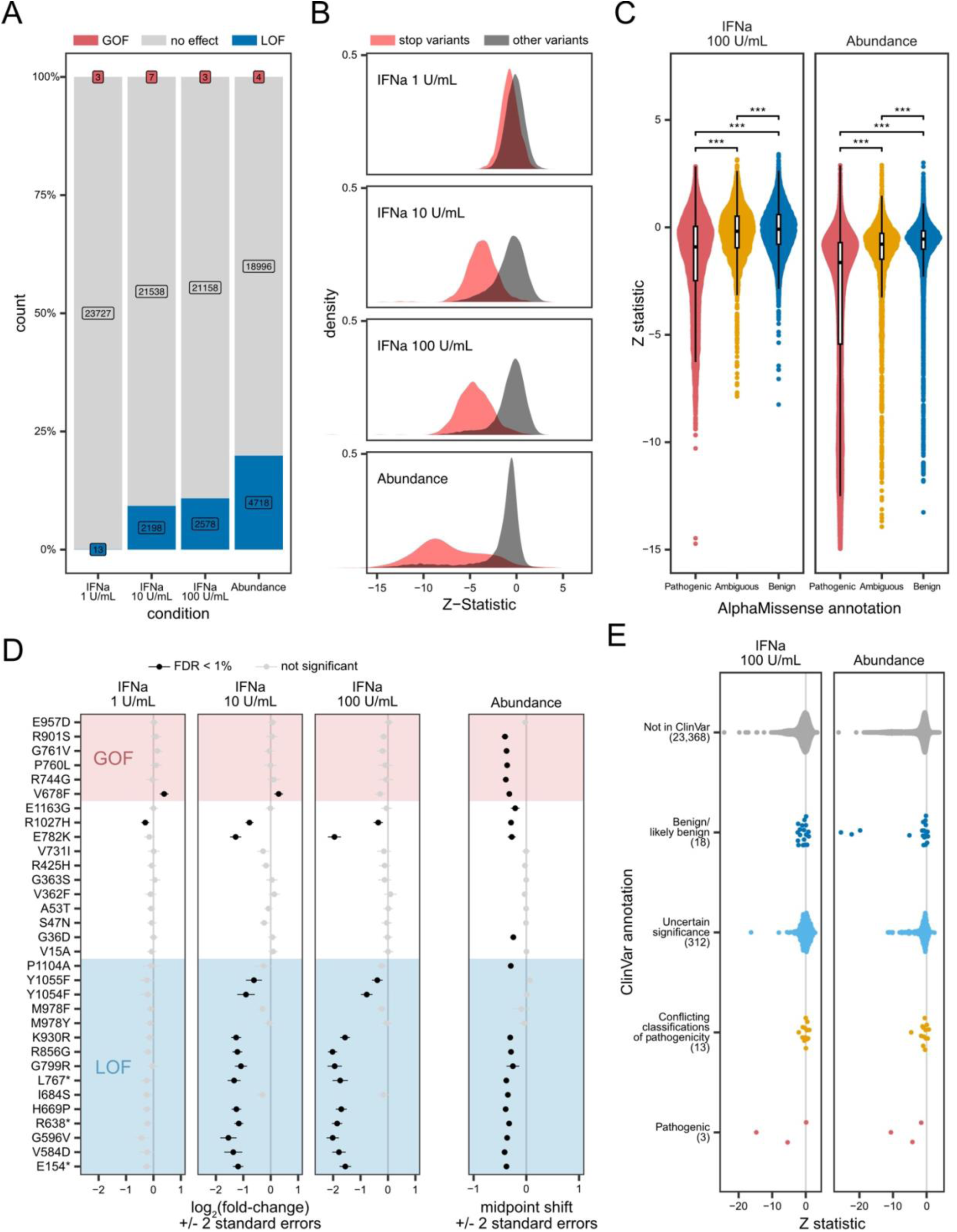
DMS results consistent with other variant effect assessments. **A**, Bar plots quantifying the frequency of LOF (blue), neutral (gray) and GOF (red) variant effects across the IFN-α and abundance DMS datasets. **B**, Histograms of stop (red) vs. nonstop (gray) variant effects, where greater separation of these two distributions indicates greater dynamic range and power to detect intermediate variant effects. The x-axis represents variant effect measured as Z-statistic of the variant normalized to WT (i.e., WT = 0), and this convention is carried through all subsequent panels. **C**, Violin plots of IFN-α (left) and abundance (right) DMS variant effects (Z-statistic, y-axis), grouped and colored by AlphaMissense annotation. **D**, DMS variant effects (log_2_(fold-change) for a panel of 32 previously characterized TYK2 variants, grouped by whether they were predicted to be gain-of-function (GOF, top), neutral (middle), or loss-of-function (LOF, bottom). **E**, Violin plots of IFN-α (left) and abundance (right) DMS variant effects (Z-statistic, x-axis), grouped and colored by ClinVar annotation.

**Supplementary Fig. 4:**
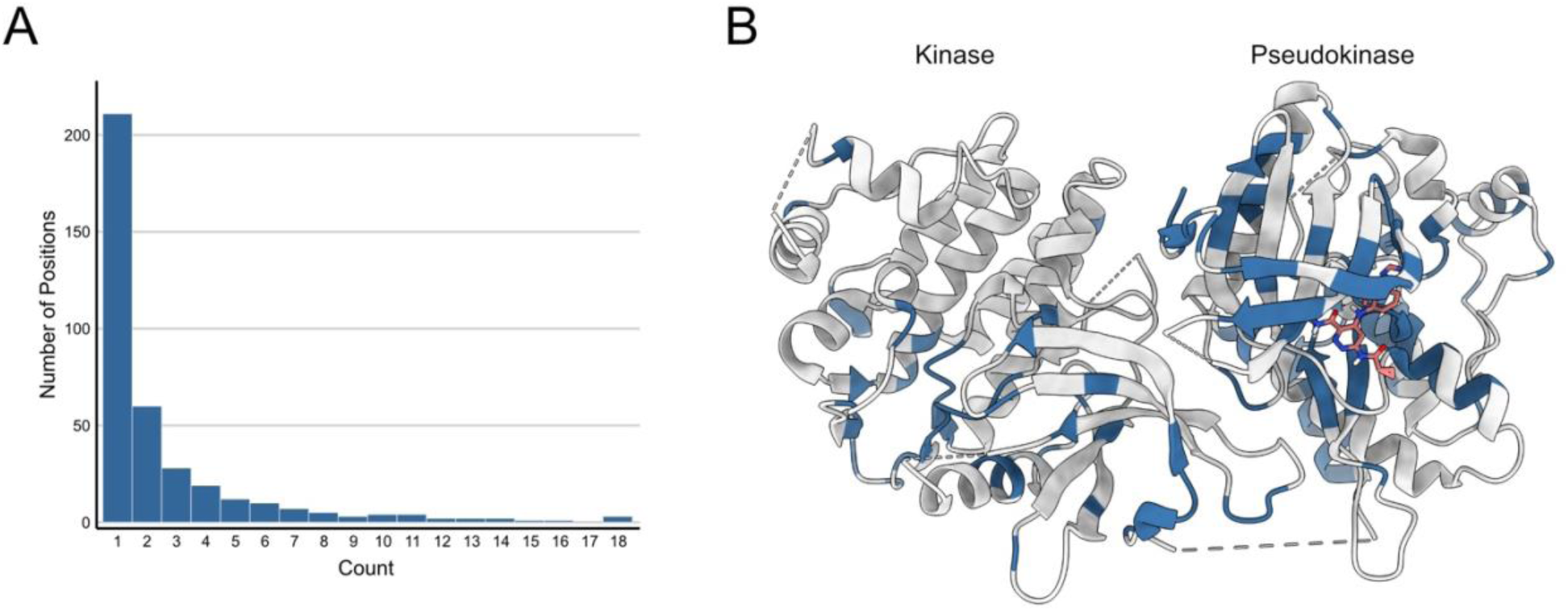
Intersecting signaling and abundance DMS data identifies allosteric and other functionally important sites. **A**, Histogram quantifying the number of amino acid positions that have X signaling-only LOF variants (“count”). **B**, TYK2kinase-pseudokinase structure (PDB: 4OLI) with amino acid positions that have at least one significant signaling-only LOF variants colored blue.

**Supplementary Fig. 5:**
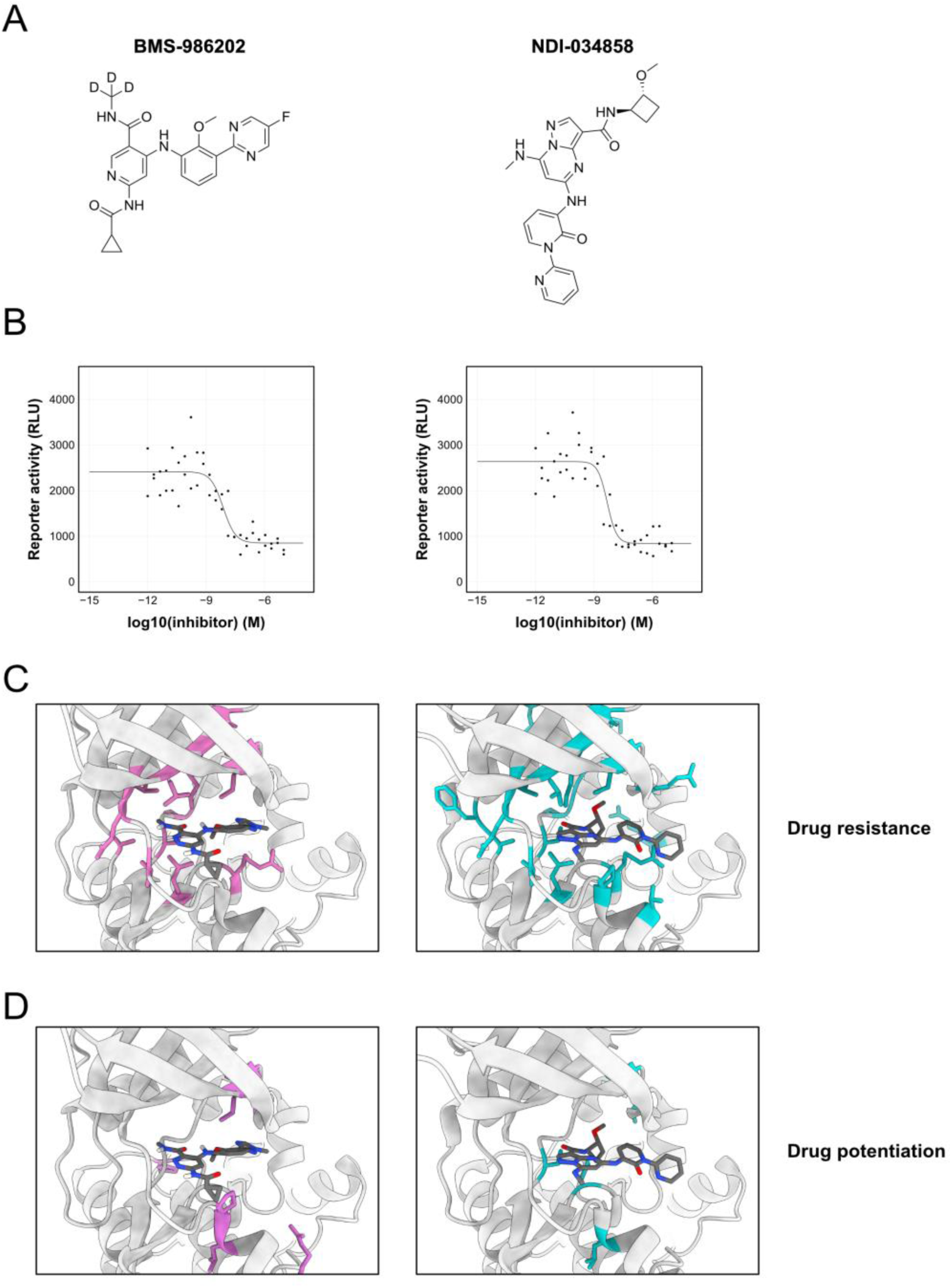
Distinct drug-resistance and drug-potentiating profiles of TYK2 allosteric inhibitors. **A**, Chemical structures of BMS-986202 (left) and NDI-034858 (right). **B**, Dose-response curves of allosteric inhibitors generated with the IFN-α reporter cell line. **C**, Structures of bound allosteric inhibitors BMS-98202 (left, PDB: 6NZP) and NDI-034858 (right, PDB: 8S9A) with residues that confer drug resistance colored pink or cyan, respectively. **D**, Structures of inhibitors with residues that confer drug-potentiation colored as in **C**.

**Supplementary Fig. 6:**
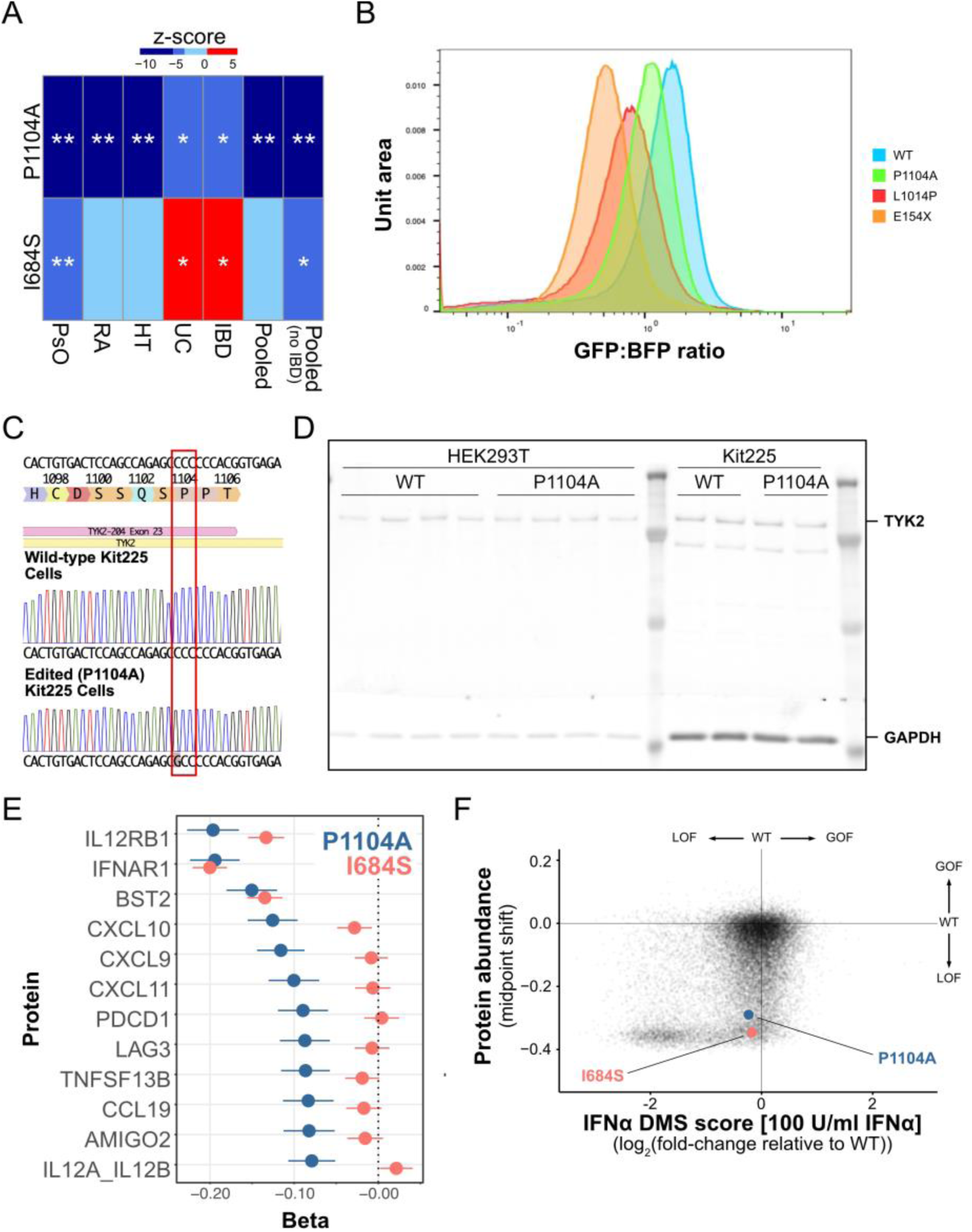
Common TYK2 variants associated with autoimmune phenotypes reduce TYK2 abundance. **A**, Effects (as z-scores) of two common partial loss-of-function missense variants of *TYK2* on relatively well-powered autoimmune traits in UKBB. (**) denotes that the variant is the most likely causal variant for the trait at the 5E-08 threshold, while (*) denotes that the variant was denoted to be the most likely causal variant for the trait at the 5E-05 significance threshold. PsO, psoriasis; RA, rheumatoid arthritis; HT, hypothyroidism; UC, ulcerative colitis; IBD, inflammatory bowel disease; Pooled, autoimmune phenotypes combined; Pooled (no IBD), same as Pooled but excluding IBD cases. **B**, Example flow cytometry data showing variants with intermediate abundance (P1104A, L1014P) relative to WT and an early stop variant (E154X). **C**, Sanger sequencing traces confirming P1104A mutation in Kit225 cells. **D**, Representative western blot of WT and P1104A TYK2 in HEK293T and Kit225 cells. **E**, Deep mutational scanning results for the IFN-α (x-axis) and protein abundance (y-axis) assays, as in **Fig. 2B**. Each point represents a unique missense or nonsense variant, with P1104A and I684S variants colored blue and pink, respectively. **F**, Effect sizes (*beta*) of association between TYK2 alleles (P1104A, blue; and I684S, pink) and abundance of twelve proteins in human plasma from UK Biobank. All twelve proteins show a significant association with. P1104A, while only the top three show a significant association with I684S. Error bars are 95% confidence intervals.

**Supplementary Fig. 7:**
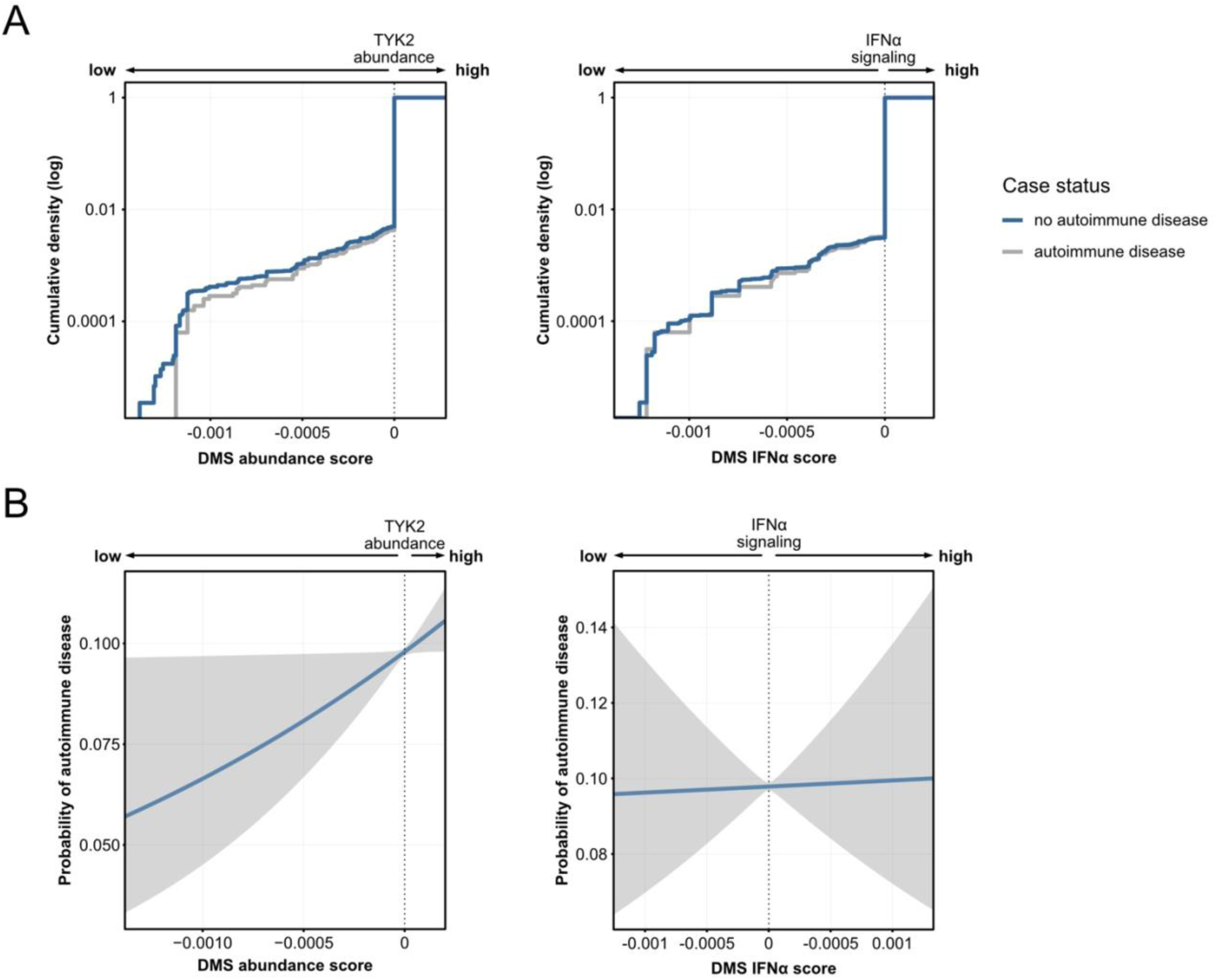
Rare variants that reduce TYK2 abundance protect against autoimmune phenotypes. **A**, Cumulative density plots quantifying the frequency of autoimmune disease in patient variants against the respective variant effect on protein abundance (left) or IFN-α signaling (right). Variants with very low abundance scores are uniquely associated with no autoimmune disease. **B**, Rare variant association test for TYK2 variants found in the UK Biobank, comparing probability of autoimmune disease (y-axis) against variant effects on TYK2 abundance (left, reproduced from main **Fig. 4E**) or IFN-α signaling (right), showing that reduced TYK2 protein abundance (left) is correlated with a reduced probability of autoimmune disease. Effects on IFN-α signaling (right) show no correlation with autoimmune disease.

**Supplementary Fig. 8:**
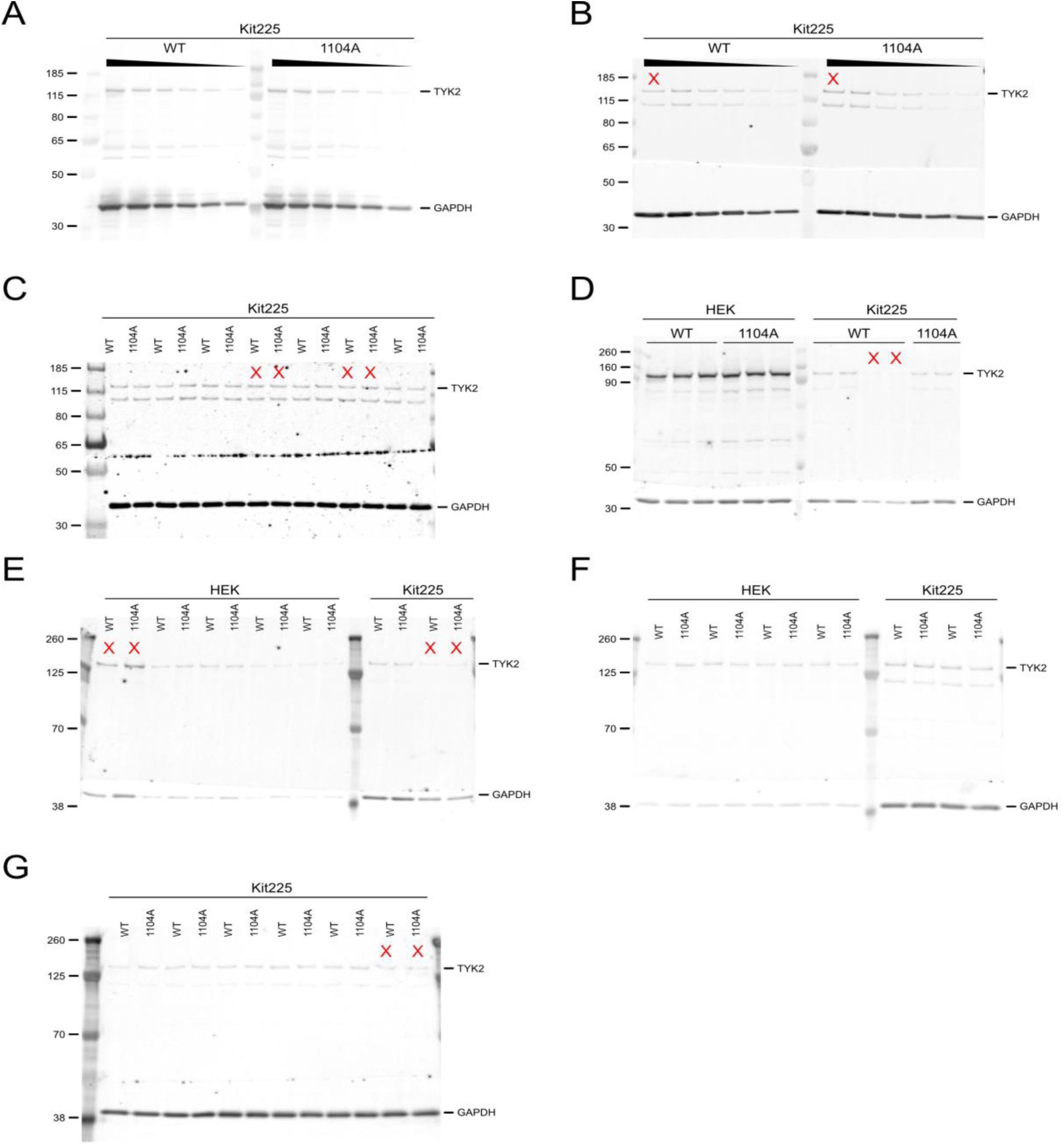
Raw western blots used to validate reduced protein abundance of P1104A. **A—G**, Images of all blots stained for TYK2 (top) and GAPDH loading control (bottom). For each lane, TYK2 band density was normalized to GAPDH band density. Lanes that were excluded are marked with a red X, determined by whether either band was outside of the linear detection range or contained blotting artifacts. Lanes are labeled by allele (WT or P1104A), with the cell background (HEK or Kit225) labelled above the alleles. See **Methods** for more details on how bands were quantified and normalized for analysis.

**Supplementary Tables**

Provided as separate Excel file.

## Notes

https://www.ncbi.nlm.nih.gov/bioproject/?term=PRJNA1291213

https://zenodo.org/records/15347448

https://github.com/octantbio/bms-dms

### Summary of Updates

Several individuals who had previously been included in the acknowledgements section were added as authors, and the conflict of interest statement was updated to include the new authors. No scientific content was changed.

### Funding

- Bristol-Myers Squibb (United States)

## References

1. Minegishi, Y. et al. Human tyrosine kinase 2 deficiency reveals its requisite roles in multiple cytokine signals involved in innate and acquired immunity. Immunity 25, 745–755 (2006).

2. Kreins, A. Y. et al. Human TYK2 deficiency: Mycobacterial and viral infections without hyper-IgE syndrome. J. Exp. Med. 212, 1641–1662 (2015).

3. Muromoto, R., Oritani, K. & Matsuda, T. Current understanding of the role of tyrosine kinase 2 signaling in immune responses. World J. Biol. Chem. 13, 1–14 (2022).

4. Hu, X., Li, J., Fu, M., Zhao, X. & Wang, W. The JAK/STAT signaling pathway: from bench to clinic. Signal Transduct. Target. Ther. 6, 402 (2021).

5. International Multiple Sclerosis Genetics Consortium (IMSGC) et al. Analysis of immune-related loci identifies 48 new susceptibility variants for multiple sclerosis. Nat. Genet. 45, 1353–1360 (2013).

6. Diogo, D. et al. TYK2 protein-coding variants protect against rheumatoid arthritis and autoimmunity, with no evidence of major pleiotropic effects on non-autoimmune complex traits. PLoS One 10, e0122271 (2015).

7. Genetic Analysis of Psoriasis Consortium & the Wellcome Trust Case Control Consortium 2 et al. A genome-wide association study identifies new psoriasis susceptibility loci and an interaction between HLA-C and ERAP1. Nat. Genet. 42, 985–990 (2010).

8. Dendrou, C. A. et al. Resolving TYK2 locus genotype-to-phenotype differences in autoimmunity. Sci. Transl. Med. 8, 363ra149 (2016).

9. Boisson-Dupuis, S. et al. Tuberculosis and impaired IL-23-dependent IFN-γ immunity in humans homozygous for a common TYK2 missense variant. Sci Immunol 3, (2018).

10. Ogishi, M. et al. Impaired IL-23-dependent induction of IFN-γ underlies mycobacterial disease in patients with inherited TYK2 deficiency. J. Exp. Med. 219, (2022).

11. Samuel, C., Cornman, H., Kambala, A. & Kwatra, S. G. A review on the safety of using JAK inhibitors in dermatology: Clinical and laboratory monitoring. Dermatol. Ther. (Heidelb.) 13, 729–749 (2023).

12. Papp, K. et al. Phase 2 trial of selective tyrosine kinase 2 inhibition in psoriasis. N. Engl. J. Med. 379, 1313–1321 (2018).

13. Armstrong, A. W. et al. Safety and efficacy of deucravacitinib in moderate to severe plaque psoriasis for up to 3 years: An open-label extension of randomized clinical trials: An open-label extension of randomized clinical trials. JAMA Dermatol. 161, 56–66 (2025).

14. Hoy, S. M. Deucravacitinib: First approval. Drugs 82, 1671–1679 (2022).

15. Jensen, L. T., Attfield, K. E., Feldmann, M. & Fugger, L. Allosteric TYK2 inhibition: redefining autoimmune disease therapy beyond JAK1-3 inhibitors. EBioMedicine 97, 104840 (2023).

16. Armstrong, A. W. et al. Tyrosine kinase 2 inhibition with zasocitinib (TAK-279) in psoriasis: A randomized clinical trial: A randomized clinical trial. JAMA Dermatol. 160, 1066–1074 (2024).

17. Danese, S. et al. P1001 Efficacy and safety of an oral tyrosine kinase 2 inhibitor VTX958 in moderately to severely active Crohn’s disease: a randomised, double-blind, placebo-controlled, phase 2 trial. J. Crohns. Colitis 19, i1853–i1854 (2025).

18. Efficacy and safety of deucravacitinib, an oral, selective tyrosine kinase 2 inhibitor, in patients with moderately to severely active ulcerative colitis: 12-week results from the phase 2 LATTICE-UC study. Gastroenterol. Hepatol. (N. Y.) 18, 6 (2022).

19. Fowler, D. M. & Fields, S. Deep mutational scanning: a new style of protein science. Nat. Methods 11, 801–807 (2014).

20. Starita, L. M. et al. Variant Interpretation: Functional Assays to the Rescue. Am. J. Hum. Genet. 101, 315–325 (2017).

21. Maes, S., Deploey, N., Peelman, F. & Eyckerman, S. Deep mutational scanning of proteins in mammalian cells. *Cell Rep*. Methods 3, 100641 (2023).

22. Jones, E. M. et al. Structural and functional characterization of G protein-coupled receptors with deep mutational scanning. Elife 9, (2020).

23. Howard, C. J. et al. High-resolution deep mutational scanning of the melanocortin-4 receptor enables target characterization for drug discovery. Elife 13, (2025).

24. Levy, D. E., Kessler, D. S., Pine, R., Reich, N. & Darnell, J. E., Jr. Interferon-induced nuclear factors that bind a shared promoter element correlate with positive and negative transcriptional control. Genes Dev. 2, 383–393 (1988).

25. Matreyek, K. A. et al. Multiplex assessment of protein variant abundance by massively parallel sequencing. Nat. Genet. 50, 874–882 (2018).

26. Cheng, J. et al. Accurate proteome-wide missense variant effect prediction with AlphaMissense. Science 381, eadg7492 (2023).

27. Sanda, T. et al. TYK2-STAT1-BCL2 pathway dependence in T-cell acute lymphoblastic leukemia. Cancer Discov. 3, 564–577 (2013).

28. Dyment, D. A. et al. Exome sequencing identifies a novel multiple sclerosis susceptibility variant in the TYK2 gene. Neurology 79, 406–411 (2012).

29. Kerner, G. et al. Homozygosity for TYK2 P1104A underlies tuberculosis in about 1% of patients in a cohort of European ancestry. Proc. Natl. Acad. Sci. U. S. A. 116, 10430–10434 (2019).

30. Yeh, T. C., Dondi, E., Uze, G. & Pellegrini, S. A dual role for the kinase-like domain of the tyrosine kinase Tyk2 in interferon-alpha signaling. Proc. Natl. Acad. Sci. U. S. A. 97, 8991–8996 (2000).

31. Staerk, J., Kallin, A., Demoulin, J.-B., Vainchenker, W. & Constantinescu, S. N. JAK1 and Tyk2 activation by the homologous polycythemia vera JAK2 V617F mutation: cross-talk with IGF1 receptor: Cross-talk with igf1 receptor. J. Biol. Chem. 280, 41893–41899 (2005).

32. Li, Z. et al. Two rare disease-associated Tyk2 variants are catalytically impaired but signaling competent. J. Immunol. 190, 2335–2344 (2013).

33. Lupardus, P. J. et al. Structure of the pseudokinase-kinase domains from protein kinase TYK2 reveals a mechanism for Janus kinase (JAK) autoinhibition. Proc. Natl. Acad. Sci. U. S. A. 111, 8025–8030 (2014).

34. Waanders, E. et al. Germline activating TYK2 mutations in pediatric patients with two primary acute lymphoblastic leukemia occurrences. Leukemia 31, 821–828 (2017).

35. Shaw, M. H. et al. A natural mutation in the Tyk2 pseudokinase domain underlies altered susceptibility of B10.Q/J mice to infection and autoimmunity. Proc. Natl. Acad. Sci. U. S. A. 100, 11594–11599 (2003).

36. Wu, P., Chen, S., Wu, B., Chen, J. & Lv, G. A TYK2 gene mutation c.2395G>A leads to TYK2 deficiency: A case report and literature review. Front. Pediatr. 8, 253 (2020).

37. Gauzzi, M. C. et al. Interferon-alpha-dependent activation of Tyk2 requires phosphorylation of positive regulatory tyrosines by another kinase. J. Biol. Chem. 271, 20494–20500 (1996).

38. Landrum, M. J. et al. ClinVar: updates to support classifications of both germline and somatic variants. Nucleic Acids Res. 53, D1313–D1321 (2025).

39. Weng, C., Faure, A. J., Escobedo, A. & Lehner, B. The energetic and allosteric landscape for KRAS inhibition. Nature 626, 643–652 (2024).

40. Philips, R. L. et al. The JAK-STAT pathway at 30: Much learned, much more to do. Cell 185, 3857–3876 (2022).

41. Glassman, C. R. et al. Structure of a Janus kinase cytokine receptor complex reveals the basis for dimeric activation. Science 376, 163–169 (2022).

42. Lé, A. M., Puig, L. & Torres, T. Deucravacitinib for the treatment of psoriatic disease. Am. J. Clin. Dermatol. 23, 813–822 (2022).

43. Tokarski, J. S. et al. Tyrosine Kinase 2-mediated Signal Transduction in T Lymphocytes Is Blocked by Pharmacological Stabilization of Its Pseudokinase Domain. J. Biol. Chem. 290, 11061–11074 (2015).

44. Chimalakonda, A. et al. Selectivity profile of the tyrosine kinase 2 inhibitor deucravacitinib compared with Janus kinase 1/2/3 inhibitors. Dermatol. Ther. (Heidelb.) 11, 1763–1776 (2021).

45. Roskoski, R., Jr. Deucravacitinb is an allosteric TYK2 protein kinase inhibitor FDA-approved for the treatment of psoriasis. Pharmacol. Res. 106642 (2023).

46. Leit, S. et al. Discovery of a potent and selective tyrosine kinase 2 inhibitor: TAK-279. J. Med. Chem. 66, 10473–10496 (2023).

47. Wallweber, H. J. A., Tam, C., Franke, Y., Starovasnik, M. A. & Lupardus, P. J. Structural basis of recognition of interferon-α receptor by tyrosine kinase 2. Nat. Struct. Mol. Biol. 21, 443–448 (2014).

48. Pines, G., Fankhauser, R. G. & Eckert, C. A. Predicting drug resistance using deep mutational scanning. Molecules 25, 2265 (2020).

49. Persky, N. S. et al. Defining the landscape of ATP-competitive inhibitor resistance residues in protein kinases. Nat. Struct. Mol. Biol. 27, 92–104 (2020).

50. Procko, E. Deep mutagenesis in the study of COVID-19: a technical overview for the proteomics community. Expert Rev. Proteomics 17, 633–638 (2020).

51. Chardon, F. M. et al. A multiplex, prime editing framework for identifying drug resistance variants at scale. bioRxiv (2023) doi:10.1101/2023.07.27.550902.

52. Estevam, G. O. et al. Mapping kinase domain resistance mechanisms for the MET receptor tyrosine kinase via deep mutational scanning. Elife 13, (2025).

53. Liu, C. et al. Discovery of BMS-986202: A clinical Tyk2 inhibitor that binds to Tyk2 JH2. J. Med. Chem. 64, 677–694 (2021).

54. Wrobleski, S. T. et al. Highly selective inhibition of tyrosine kinase 2 (TYK2) for the treatment of autoimmune diseases: Discovery of the allosteric inhibitor BMS-986165. J. Med. Chem. 62, 8973–8995 (2019).

55. Ban, M. et al. Replication analysis identifies TYK2 as a multiple sclerosis susceptibility factor. Eur. J. Hum. Genet. 17, 1309–1313 (2009).

56. Sun, B. B. et al. Plasma proteomic associations with genetics and health in the UK Biobank. Nature 622, 329–338 (2023).

57. Plenge, R. M., Scolnick, E. M. & Altshuler, D. Validating therapeutic targets through human genetics. Nat. Rev. Drug Discov. 12, 581–594 (2013).

58. Schindelin, J. et al. Fiji: an open-source platform for biological-image analysis. Nat. Methods 9, 676–682 (2012).

59. Landrum, M. J. et al. ClinVar: public archive of relationships among sequence variation and human phenotype. Nucleic Acids Res. 42, D980–5 (2014).

60. Pettersen, E. F. et al. UCSF ChimeraX: Structure visualization for researchers, educators, and developers. Protein Sci. 30, 70–82 (2021).

61. Kurki, M. I. et al. FinnGen provides genetic insights from a well-phenotyped isolated population. Nature 613, 508–518 (2023).

62. Bycroft, C. et al. The UK Biobank resource with deep phenotyping and genomic data. Nature 562, 203–209 (2018).

63. Langefeld, C. D. et al. Transancestral mapping and genetic load in systemic lupus erythematosus. Nat. Commun. 8, 16021 (2017).

64. Tsoi, L. C. et al. Identification of 15 new psoriasis susceptibility loci highlights the role of innate immunity. Nat. Genet. 44, 1341–1348 (2012).

65. Okada, Y. et al. Genetics of rheumatoid arthritis contributes to biology and drug discovery. Nature 506, 376–381 (2014).

66. Liu, J. Z. et al. Association analyses identify 38 susceptibility loci for inflammatory bowel disease and highlight shared genetic risk across populations. Nat. Genet. 47, 979–986 (2015).

67. International Multiple Sclerosis Genetics Consortium. Multiple sclerosis genomic map implicates peripheral immune cells and microglia in susceptibility. Science 365, eaav7188 (2019).

68. López-Isac, E. et al. Influence of TYK2 in systemic sclerosis susceptibility: a new locus in the IL-12 pathway. Ann. Rheum. Dis. 75, 1521–1526 (2016).

69. Betz, R. C. et al. Genome-wide meta-analysis in alopecia areata resolves HLA associations and reveals two new susceptibility loci. Nat. Commun. 6, 5966 (2015).

70. Mbatchou, J. et al. Computationally efficient whole-genome regression for quantitative and binary traits. Nat. Genet. 53, 1097–1103 (2021).

71. Yang, J., Lee, S. H., Goddard, M. E. & Visscher, P. M. GCTA: a tool for genome-wide complex trait analysis. Am. J. Hum. Genet. 88, 76–82 (2011).

72. Wang, G., Sarkar, A., Carbonetto, P. & Stephens, M. A simple new approach to variable selection in regression, with application to genetic fine mapping. J. R. Stat. Soc. Series B Stat. Methodol. 82, 1273–1300 (2020).

73. Backman, J. D. et al. Exome sequencing and analysis of 454,787 UK Biobank participants. Nature 599, 628–634 (2021).

74. McLaren, W. et al. The Ensembl Variant Effect Predictor. Genome Biol. 17, 122 (2016).

75. Lee, S. et al. Optimal unified approach for rare-variant association testing with application to small-sample case-control whole-exome sequencing studies. Am. J. Hum. Genet. 91, 224–237 (2012).

76. McCaw, Z. R. et al. An allelic-series rare-variant association test for candidate-gene discovery. Am. J. Hum. Genet. 110, 1330–1342 (2023).

77. Wang, X. et al. CRISPR-DAV: CRISPR NGS data analysis and visualization pipeline. Bioinformatics 33, 3811–3812 (2017).

